# Whole embryo barcoding reveals convergent origins of mesodermal cells during mammalian gastrulation

**DOI:** 10.64898/2026.11.11.765433

**Authors:** Marine A. Secchi, Peter J. Martin, Jake Cornwall-Scoones, Debra van Egeren, Peyton J. Dooley, Erica Y. Liu, Nianping Liu, Anastasiia Masaltseva, Hunter D. King, Rayyan T. Jokhai, Xiaojie Qiu, Kyle M. Loh, Berthold Göttgens, Fernando D Camargo, Sarah J. Bowling

**Affiliations:** Department of Developmental Biology, Stanford University School of Medicine, Stanford, CA, USA; Department of Cell and Developmental Biology (CDB), University College London, Gower Street, London WC1E 6BT, UK; The Gurdon Institute, University of Cambridge, Tennis Court Road, Cambridge CB2 1QN, UK; Stem Cell Program, Boston Children’s Hospital, Boston, MA, USA; Department of Stem Cell and Regenerative Biology, Harvard University, Cambridge, MA, USA; Stanford Cancer Institute, Stanford University School of Medicine, Stanford, CA, USA; Department of Genetics, Stanford University School of Medicine, Stanford, CA, USA; Basic Sciences and Engineering Initiative, Betty Irene Moore Children’s Heart Center, Lucile Packard Children’s Hospital, Stanford, CA, USA; Department of Computer Science, Stanford University, Stanford, CA, USA; Institute for Stem Cell Biology and Regenerative Medicine, Stanford University School of Medicine, Stanford, CA, USA; Bioengineering Department, Stanford University, Stanford, CA, USA; Cambridge Stem Cell Institute, Cambridge, UK; Department of Hematology, University of Cambridge, Cambridge, UK

**Author notes:** Contributed equally (Peter J. Martin, Jake Cornwall-Scoones).

## Abstract

Classical fate-mapping and recent single-cell trajectory studies have suggested that mesoderm diversification during gastrulation proceeds through progressive fate bias and branching lineage segregation. Here, we combine inducible CRISPR-based lineage recording with single-cell RNA sequencing to reconstruct clonal relationships across mouse embryos barcoded at primitive streak onset and analyzed at early organogenesis. Mesodermal clones diverged after streak formation, but their developmental outputs remained extensively overlapping, revealing a clonal architecture that is not a simple branching lineage tree. Using sampling-aware statistical models, we quantify distinct clonal origins of paraxial mesoderm across the anterior-posterior axis, show that endothelial cells arise from multiple mesoderm origins but demonstrate transcriptional differences that correspond to anatomical position and clonal lineage, and identify clonal sharing between extra-embryonic mesoderm and embryonic derivatives, including cardiac lineages and a shared hindlimb-allantois progenitor. In the heart, progenitors corresponding to classical heart fields remain multipotent, contributing both myocardial and non-myocardial cell types. Together, our findings support a model in which mesodermal diversity is assembled through spatially distinct, fate-biased progenitor pools whose descendants converge on shared cell states. Our findings challenge simple branching models of mesoderm diversification and provide a clonal framework for understanding how tissue diversity emerges in the embryo.

## Introduction

A central question in developmental biology is how embryonic progenitors diversify to generate the full range of tissues in the body. In the mammalian embryo, gastrulation marks the transition from pluripotent epiblast cells to spatially organized germ layers which lay the foundation for future organ progenitors to emerge^1^. Classical fate-mapping studies in the mouse using tissue grafting, dye labelling, recombinases, and retrospective clonal analysis, established many of the foundational principles of this process^2–6^. These studies showed that progenitors are regionally organized within the epiblast and primitive streak, and that mesodermal derivatives are allocated in a broadly ordered anterior-posterior sequence as cells ingress through the streak. However, these studies only traced one progenitor population at a time and therefore cannot reveal how multiple cell types arise in parallel from shared or distinct clonal origins. Furthermore, classical approaches largely rely on embryo culture with imaging readouts which constrains fate mapping to short developmental windows and provides no molecular information about the cells being traced. As a result, it remains unclear whether the classical model of graded allocation and progressive branching fate restriction fully explains how mesodermal diversity is generated during mammalian gastrulation.

Recent single-cell transcriptomic studies have provided high-resolution maps of cell states during gastrulation^7–11^. These studies have greatly refined our understanding of mesodermal differentiation and have revealed continuous transitions between emerging progenitor states that are broadly consistent with progressive lineage commitment. However, transcriptional similarity does not necessarily reflect developmental ancestry: cells that converge on a common transcriptional identity may arise from distinct progenitors, while clonally related cells may rapidly diverge in gene expression as they enter different morphogenetic contexts. Resolving developmental relationships therefore requires lineage tracing alongside molecular characterization. Recent advances in genetic lineage recording, including inducible DNA barcoding and phylogenetic lineage reconstruction, have enabled increasingly comprehensive analyses of cellular ancestry during mammalian development^12–18^. These studies have provided important insights into the diversification of embryonic lineages and the progressive restriction of developmental fates. However, a central challenge remains to understand how these lineage relationships translate into the formation and spatial organization of differentiated tissues. In particular, the extent to which common cell identities arise through distinct lineage routes, and how clonal relationships vary across anatomical regions remain incompletely understood.

Here, we combine whole-embryo single-cell RNA sequencing with inducible CRISPR-based lineage recording using CARLIN^19^ and DARLIN^20^ models to generate a high-resolution clonal lineage atlas of mouse gastrulation. In these models, doxycycline (Dox) induction activates CRISPR editing of transcribed barcode arrays to result in heritable sequence changes that can be recovered alongside gene expression from individual cells. Cells carrying the same barcode can be identified as members of a clone, while their transcriptomes reveal the cell types to which that clone contributed. DARLIN extends CARLIN by using a fused Cas9-terminal deoxynucleotidyl transferase to increase insertion events following barcode editing which increase barcode diversity. Importantly, we use these models to trace cell fate during development, which we define as the developmental destination of a cell in its native, undisturbed environment^21^. Throughout the manuscript the term “progenitor” is used to define cells that are preferentially fated, though not necessarily committed or specified, to give rise to a particular cell type.

By barcoding embryos at the onset of primitive streak formation and sampling multiple regions from the embryo at early organogenesis, we simultaneously infer cell identity, coarse-grained anatomical location, and developmental ancestry, allowing us to measure lineage coupling between cell types across the entire embryo. We identify recurrent patterns of clonal contribution that reveal how mesodermal lineages diversify during gastrulation and subsequently contribute to distinct anatomical regions and differentiated cell types. In particular, our data reveal that lineage convergence, whereby progenitors diverge in fate yet retain overlapping developmental outputs, is a prominent feature of tissue formation, meaning that transcriptionally similar cell types can arise through distinct lineage routes. We identify lineage convergence in the formation of somitic mesoderm, endothelium, allantois, and cardiomyocytes, and uncover lineage relationships between the embryonic and extra-embryonic mesoderm, including preferential clonal coupling between the allantois and hindlimb mesoderm, and shared clones between the yolk sac and cardiomyocytes. Together, our findings reveal how overlapping progenitor contributions and spatially organized clonal relationships shape the formation of embryonic tissues, and provide a framework for understanding how developmental ancestry relates to cell identity and anatomical organization during mammalian development.

## Results

### High-resolution clonal tracing of whole barcoded mouse embryos

We set out to barcode embryos as early fate restrictions emerge in the epiblast, just prior to and coinciding with the onset of gastrulation. We collected seven barcoded embryos at embryonic day (E)9.5 (Figure S1A; somite counts ranging 16-22 somite pairs): two induced at E5.5 (iE5.5), and five induced at E6.5 (iE6.5). Each embryo was dissected into head, trunk (trunk + heart for embryo 1), tail, and visceral yolk sac (YS) prior to scRNA-seq (Figure 1A,S1B), which improved our ability to confidently annotate cells (Figure 1B, Methods, Figure S2), and provided us with coarse-grained anatomical information for the barcode analysis. In total, our dataset contains 131,447 cells, from which we identified 37 distinct cell types across multiple different regions and germ layers (Figure S2, Table S1).

**Figure 1:**
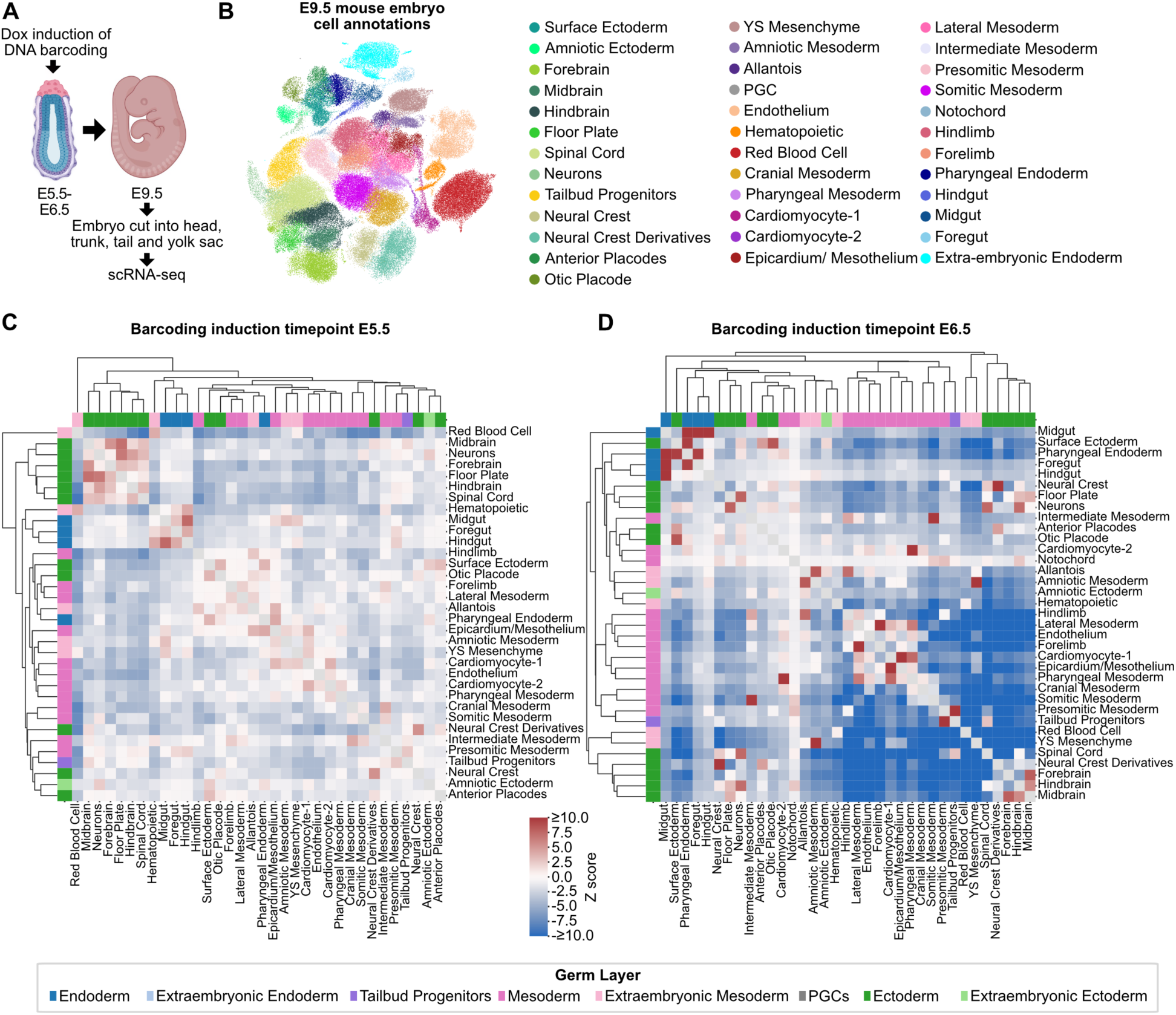
High-resolution tracing of the early post-implantation mammalian embryo. (A) Schematic of experimental design. (B) UMAP of E9.5 cells colored by cell-type annotation. (C,D) Hierarchically clustered dendrograms of cell types based on correlations between lineage-coupling Z-scores for barcodes induced at E5.5 (C) or E6.5 (D). Cell types are colored according to germ-layer annotation. Only clones containing ≥3 recovered cells are included.

To define the window of barcode labelling following Dox induction in CARLIN embryos, we collected embryos at 8, 24, 36, 48, and 60 h after induction on the sixth day of pregnancy (E6) and measured editing rates over time (Figure S3A). Editing was detectable by 8h (∼7% of CARLIN transcripts were edited), increased to an average of 29% editing by 24h across two litters, and plateaued by 36h across four litters collected at 36-60h (Figure S3B). These data indicate that the majority of barcoding occurs between 8 and 36 h following Dox induction. This is similar to the editing kinetics in the DARLIN system which we have previously demonstrated occurs over ∼24h^20^. In the absence of Dox, editing was negligible in both the CARLIN (<0.2%) and DARLIN models^20^, consistent with minimal background labelling. To relate this labelling window to developmental stage, we collected DARLIN embryos at E6.5, E7.0, and E7.5, assessed their morphology and Brachyury expression, and staged them using published criteria^22^. These analyses indicate that Dox induction at E6.5 labels cells from approximately the early primitive streak to late streak/early headfold stages (Figure S3C,D).

For the majority of cell types, we obtained a CARLIN or DARLIN transcript from ∼70% cells in our dataset across embryos (Figure S4A, Table S1). An exception is red blood cells, which have lower capture efficiencies averaging ∼30%. While editing rates were generally uniform across cell types, we observed almost negligible labelling in the extra-embryonic endoderm. Barcode editing rates for all other cell types ranged 21-55% for CARLIN embryos and 49-73% for DARLIN embryos (Figure S4B, Table S1). Following removal of barcodes that did not meet our strict filtering thresholds (Methods; Figure S3E-G), an average of 15% of cells in our dataset contained useful barcode information. To assess whether barcode recovery varied systematically between cell types, we modelled the observed variation using a beta-binomial distribution, which accounts for overdispersion beyond that expected from simple random sampling (Methods; Figure S5A,B). This model explained much of the variation in barcode recovery across cell types, although certain populations, including red blood cells, extraembryonic endoderm, and amniotic ectoderm, deviated from the expected distribution. Based on these findings, we assumed equivalent barcode recovery rates across cell types in subsequent analyses, and excluded outlier populations from certain statistical tests. Finally, assuming an E9.5 embryo size of approximately 500,000 cells, we estimated that cells with usable barcodes represented 0.2-1.2% of the total cell population per embryo (Figure S5C).

First, we investigated the fate patterns exhibited by cells at our two barcoding induction time points by performing lineage coupling analysis^23^ (Methods). Here, lineage coupling between cell states was quantified using a permutation-based clonal sharing statistic: positive z-scores indicate enriched clonal sharing and fate bias of clones towards the cell states, whereas negative values indicate reduced sharing relative to a null model lacking fate bias, consistent with clonal divergence. This method accounts for biases introduced by the sampling and by the differential abundance of cell types in the E9.5 embryo. Following barcode induction at E5.5, cell states clustered largely according to germ layer identity (Figure 1C), with modules corresponding to neural ectoderm, mesodermal derivatives, and endodermal tissues. Given previous work^1–5^, we interpret this as supporting regionalized fate within the pre-gastrulation epiblast as opposed to demonstrating that cell commitment or specification has occurred by this stage. Induction at E6.5 produced higher Z-scores, consistent with increased bias within clones and progressive lineage restriction following gastrulation (Figure 1D). Lineage-coupling Z-scores were positively correlated between each individual E6.5 embryo and the remaining embryos pooled (Figure S6,7; ρ=0.64-0.78 across 465 cell-type pairs), supporting reproducibility of the broad coupling structure across embryos. Individual coupling scores nevertheless varied (Figure S7), therefore we performed embryo-wise assessment for each of the specific lineage relationships discussed below.

### Mapping the diversification of mesoderm lineages

We next asked whether clones share recurrent patterns of cell-type contribution, which we term “lineage routes”, that could reflect common behaviors of similar progenitor populations at the time of labelling. To identify these patterns, we fitted a Dirichlet-multinomial mixture (DMM) model to the recovered cell-type counts of each clone (Methods)^24,25^. This model identifies recurrent contribution profiles, which we call ‘modes’, and estimates the probability that each clone belongs to each mode. The DMM framework is well suited to sparse clonal data as it accounts for differences in recovered clone size, allows clones with related behaviors to vary in their precise cell type composition, and explicitly models the sampling process of the assay. Model comparison favored eight modes that represent distinct cell-type contribution profiles that best described the data (Figure S8A). Across embryos, ∼83-90% clones were assigned to a mode (odds ratio threshold of 2 for the best-fitting mode vs the next), and all eight modes were represented in every embryo, although their relative frequences varied (Figure S8B). Mode contribution profiles were largely stable when the minimum recovered clone size was increased and individual embryos were omitted from fitting, although some profiles showed embryo-dependent variation (Figure S8C-E). Modes 2-8 showed preferential contribution to either: 1) neurectoderm derivatives (brain, neural crest, spinal cord, Mode 2); 2) extra-embryonic mesoderm (Mode 3); 3) lateral and pharyngeal mesoderm (Mode 4); 4) somitic and cranial mesoderm (Mode 5); 5) hindlimb mesoderm (Mode 6); 6) posterior ectoderm and mesoderm derivatives (tailbud progenitors, spinal cord, somitic mesoderm; Mode 7); 7) endoderm derivatives and surface ectoderm (Mode 8; Figure 2A-D; S8F; S9A). We interpret these seven recurrent patterns as representing seven distinct lineage routes within the embryo. Mode 1 was characterized by single-lineage clones, mostly within the red blood cells, YS mesenchyme, cranial mesoderm, and forebrain, which could reflect early restriction and/ or increased clonal expansion of these cell types (Figure S9). Together, these lineage routes reveal distinct patterns of mesodermal lineage contribution following E6.5 induction and largely recapitulate previously described clonal relationships in the embryo^1–5^.

**Figure 2:**
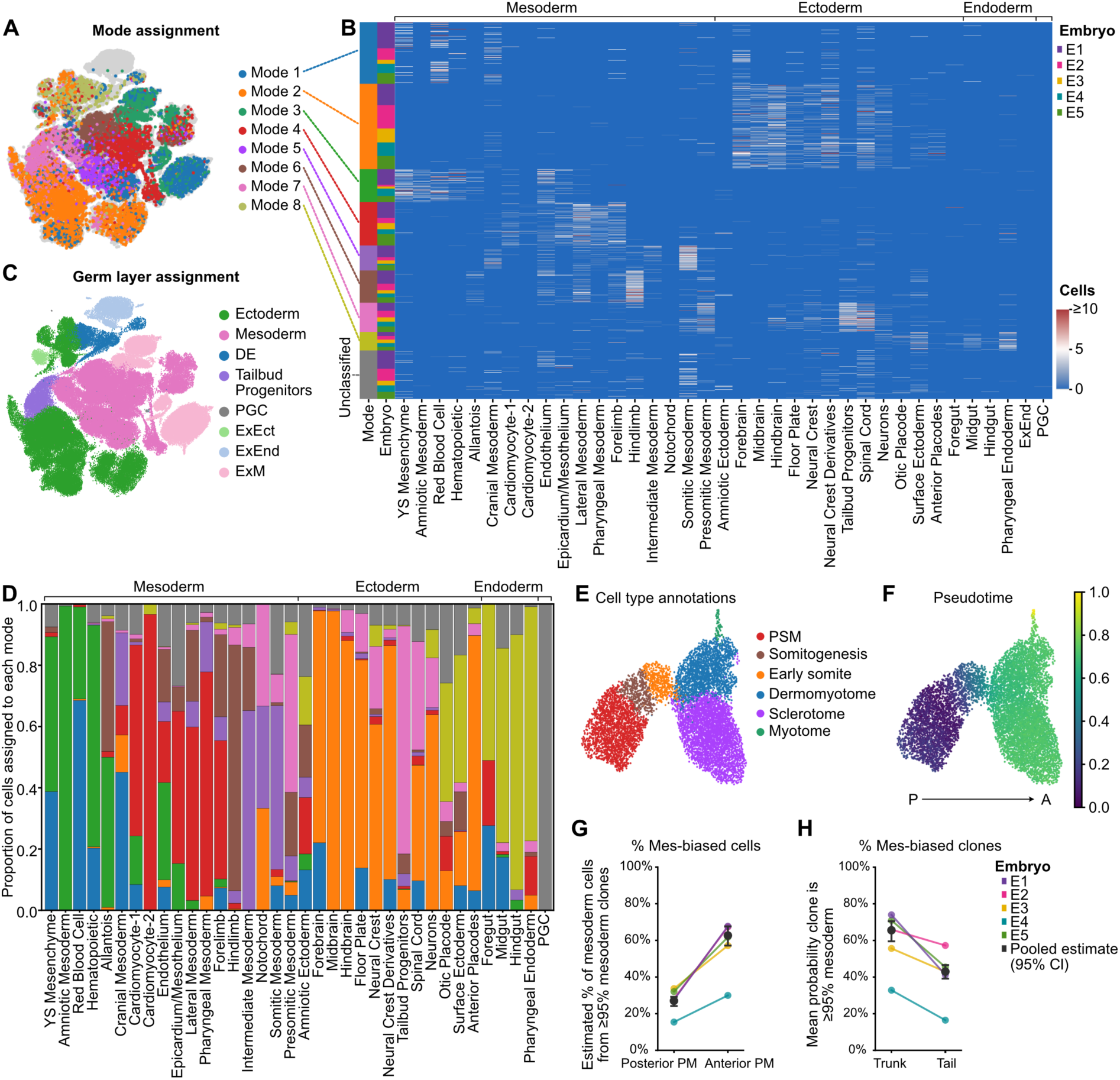
Diversification of mesoderm lineages. (A) UMAP colored according to the Dirichlet-multinomial mixture (DMM) mode assigned to each cell’s clone. Clones were assigned to a mode if its posterior probability was at least twice that of the next most probable mode; cells without a mode assignment (Unclassified) are not shown. (B) Heatmap showing the cell-type composition of individual clones, grouped by DMM mode and embryo. Rows represent unique barcodes and columns represent cell types, grouped by germ layer. Color indicates the number of recovered cells, capped at 10. Only clones containing ≥3 recovered cells are included. (C) UMAP colored by germ-layer annotation, illustrating the correspondence between DMM mode assignments and germ-layer identity. (D) Distribution of DMM modes across cell types, grouped by germ layer. Stacked bars indicate the proportion of cells assigned to each mode, with unclassified cells shown in grey. Only clones containing ≥3 recovered cells are included. (E) UMAP of presomitic and somitic mesoderm cells following subclustering, Leiden clustering and reannotation. (F) Diffusion pseudotime (DPT) scores projected onto the paraxial mesoderm UMAP. (G,H) Model-estimated probability that a sampled paraxial mesoderm cell belongs to a strongly mesoderm-biased clone (≥95% mesoderm), comparing posterior and anterior paraxial mesoderm populations (G) and trunk and tail libraries (H) using the sampling-aware generative model M4. Points represent individual embryos. All panels show data from embryos barcoded at E6.5 (iE6.5).

Given the broad range of lineage routes identified within the emerging mesoderm (Figure 2D), we focus the remainder of this study on mesoderm diversification.

We noted that the somitic mesoderm contained clones observed in two modes: one dominated by a range of mesoderm cell types including intermediate, lateral, and cranial mesoderm (mode 5); and another containing presomitic mesoderm, tailbud progenitors, and ectodermal derivatives including spinal cord and neural crest (mode 7). Paraxial mesoderm (comprising somitic and presomitic mesoderm) generates the somites, which give rise to axial skeletal and muscle tissues. Previous lineage inference and fate-mapping studies support distinct developmental routes to anterior and posterior somites, including a direct primitive streak origin for anterior paraxial mesoderm and a neuromesodermal progenitor (NMP) contribution to posterior paraxial mesoderm^6,26,27^. Our observations are consistent with these dual origins. We next asked whether the two classes of somitic mesoderm clones contribute to distinct paraxial mesoderm differentiation states and axial identities.

We first investigated whether the two clonal profiles observed within the somitic mesoderm reflect likely differences in underlying lineage composition rather than incomplete recovery of ectodermal descendants caused by under-sampling of the embryo. For this, we subset the data to clones containing at least one paraxial mesoderm cell, and fit four generative models that incorporate population clone size, mesoderm composition, and embryo-specific cell recovery (Figure S10A; Methods). The best-performing model across held-out embryos allowed for both a mesoderm-only clone population and variable lineage composition among mixed clones that contain both mesodermal and non-mesodermal descendants (Figure S10B-D). Under this model, an estimated 52% of mesoderm cells within the trunk and tail libraries belonged to clones comprising at least 95% mesoderm cells (hereafter referred to as Mes-biased), supporting the existence of a substantial predominantly mesodermal population (Figure S11A). The estimated proportion of Mes-biased cells was largely stable across embryos with the exception of Embryo 4. Notably, this embryo was also the developmentally “youngest” collected based on somite number (16 somite pairs in E4 versus 20-22 in all other embryos), raising the possibility that earlier labelling of progenitors results in more germ layer sharing (Figure S1). Regardless, across all embryos, incomplete recovery of ectodermal descendants from the mesoderm-containing clones was insufficient to explain the observed clonal composition patterns within the somitic mesoderm.

Having established evidence for predominantly mesodermal and mixed clonal contributions, we next investigated which non-mesodermal cell types were associated with the inferred mixed clones. Specifically, we estimated the probability of detecting different non-mesodermal cell types among sampled clones that were predicted to contain less than 95% mesoderm. Within these mixed clones, on average 60% of non-mesodermal cells were inferred to be ectodermal, ∼35% were inferred to be tailbud progenitors, while 4% were inferred to be endodermal cells (Figure S11B). Given that the tailbud is a known niche for neuromesodermal progenitors (NMPs), these findings are consistent with an NMP contribution to a subset of paraxial mesoderm. We therefore investigated whether these distinct clonal contribution patterns varied along the anterior-posterior axis, with mixed clones preferentially contributing to posterior paraxial mesoderm and Mes-biased clones to anterior somitic identities. We restricted the dataset to paraxial mesoderm and its derivatives, recomputed the neighborhood graph, and inferred developmental progression using diffusion pseudotime^28^ (DPT; Figure 2E,F). Expression of established regulators of somitogenesis and *Hox* genes enabled finer cell-type annotation and confirmed that the trajectory recapitulates axial identity (Figure 2F, S11C-E). Cells belonging to Mes-biased clones were enriched at late pseudotime positions that represent more anterior regions (Figure 2G, S11F): an estimated 24% barcoded presomitic mesoderm belonged to Mes-biased clones, compared with an estimated 67% anterior myotome cells (average percentages across embryos). Accordingly, we also identified an enrichment of Mes-biased cells in the more anterior trunk libraries vs tail libraries (66% clones vs 43% clones are Mes-biased, respectively; Figure 2H). Together, these data support distinct lineage contributions along the anterior-posterior axis: Mes-biased clones preferentially contribute to anterior somitic identities while those with mesoderm-ectoderm sharing and tailbud-associated ancestry are enriched in posterior paraxial mesoderm. The clonal links to spinal cord and tailbud progenitors are consistent with an NMP contribution to posterior paraxial mesoderm that has been described previously^6,26,27^, providing us with important validation of our dataset.

### Endothelial cell heterogeneity driven by multiple distinct clonal origins

We next examined the clonal origins and transcriptional heterogeneity of endothelial cells (ECs), which line blood vessels throughout the embryo and establish the embryonic circulation. Endothelial cells receive contributions from multiple mesodermal sources including the lateral plate, extra-embryonic, and paraxial mesoderm, and recent work has demonstrated that ECs acquire tissue-specific molecular identities during embryonic development^29^. These findings raise the question of how endothelial ancestry is organized across the embryo and whether different progenitor contributions are associated with endothelial transcriptional heterogeneity. In our dataset, ECs formed a single transcriptional cluster on the UMAP marked by *Cdh5* expression, which encodes VE-cadherin (Figure 3A,B), along with *Pecam1*, *Plap*, and *Erg* (Figures S12A). Despite this shared endothelial identity, ECs were found in multiple modes (Figure 2B), consistent with the convergence of multiple distinct mesodermal lineages onto a common endothelial fate. To determine whether ECs arise locally within each region of the embryo, we leveraged the anatomical resolution of our dissection strategy to quantify clonal coupling between ECs and neighboring non-EC populations across embryonic regions. Using embryo-specific permutation tests (Methods), we observed that ECs preferentially shared barcodes with non-ECs from the same region, whereas sharing across regions was generally lower than expected under the null (Figure 3C,D). This pattern was also observed in embryo-specific analyses, although its magnitude varied (Figure S12B). These findings support regionalized endothelial ancestry, consistent with local progenitor contributions. We next asked whether finer clonal subclasses of ECs exist within tissues. We focused on four modes with the best representation of clones across all embryos which showed preferential contribution to: 1) ExM cells, 2) lateral and pharyngeal mesoderm; 3) somitic mesoderm; and 4) hindlimb mesoderm (Figure 3E,F;S12C). Together, these findings support that during gastrulation, mesodermal progenitors retain the capacity to form endothelium after otherwise diverging in fate, such that distinct regions of the embryo locally produce blood vessels.

**Figure 3:**
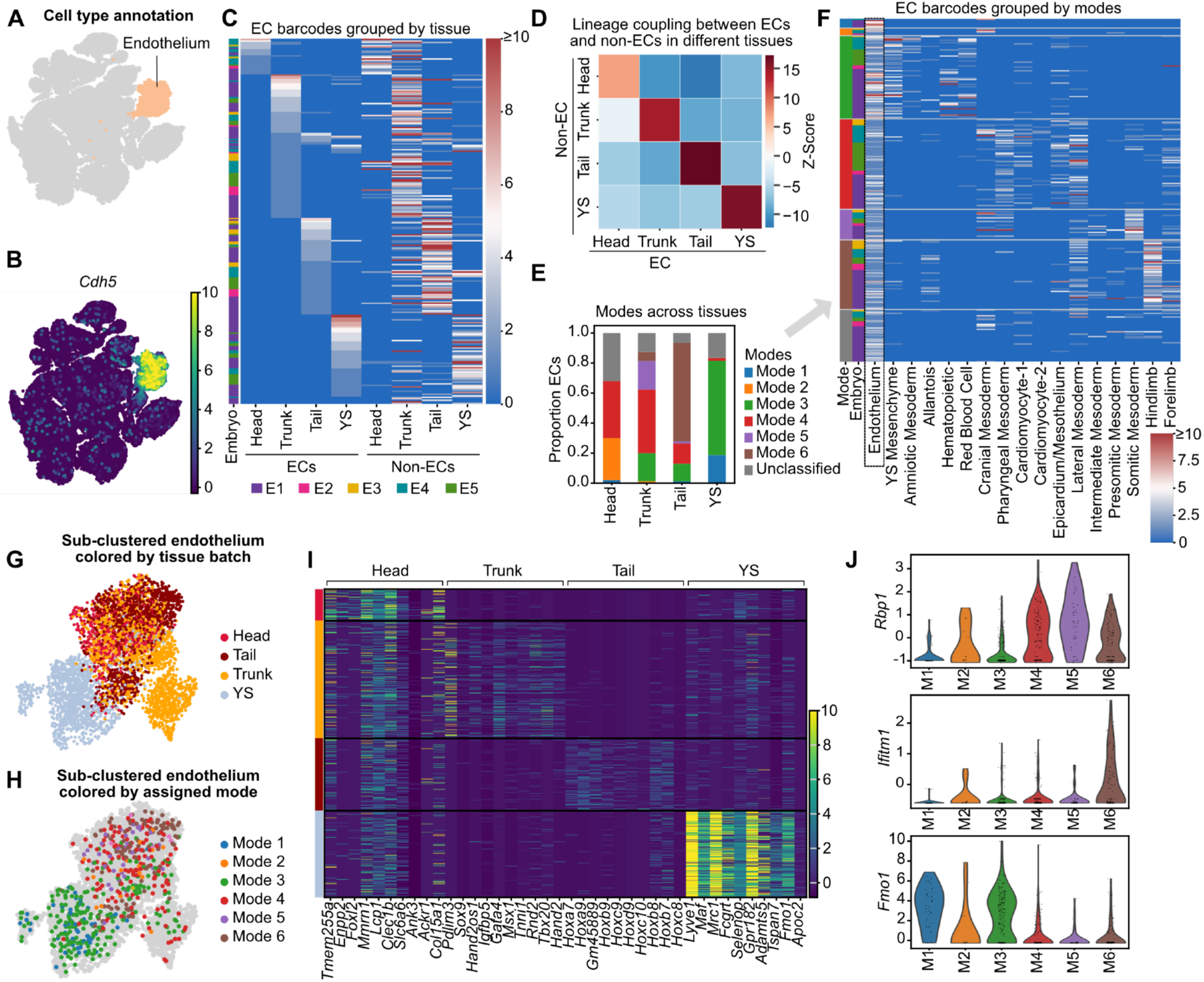
Diverse origins of endothelium associated with distinct transcriptional signatures. (A,B) UMAP of endothelial cells (ECs), colored by cell-type annotation (A) or *Cdh5* expression (B). (C) Heatmap showing the distribution of EC-containing clones across ECs and non-ECs from different anatomical regions. Rows represent unique barcodes and columns represent cell populations grouped by dissection library and embryo. Color indicates the number of recovered cells, capped at 10. Only clones containing ≥3 recovered cells are included. (D) Heatmap of conditional lineage-coupling Z-scores between ECs and non-ECs from the head, trunk, tail and yolk sac libraries. (E) Distribution of DMM mode assignments among ECs from the four dissection libraries. (F) Heatmap showing the distribution of EC-containing clones across mesoderm cell types, grouped by DMM mode assignment and embryo. Rows represent unique barcodes; color indicates the number of recovered cells, as in C. Only clones containing ≥3 recovered cells are included. (G,H) UMAP of subclustered ECs colored by dissection library (G) or assigned DMM mode (H). (I) Heatmap showing the top 10 differentially expressed genes, ranked by adjusted P-value, identified between EC groups of different tissue origins. (J) Violin plots showing log-normalized scaled expression of three differentially expressed genes identified between EC barcode groups: *Rbp1*, *Ifitm1* and *Fmo1*. All panels show data from embryos barcoded at E6.5 (iE6.5).

Given the distinct origins of ECs, we asked whether anatomical location and/or clonal origin drives EC heterogeneity. To address this, we subclustered endothelial cells and visualized them by UMAP which demonstrated some degree of clustering by both tissue of origin (Figure 3G) and clonal grouping (Figure 3H). Differential gene expression (DGE) on pseudo-bulked ECs from different areas of the embryo revealed a large number of differentially expressed genes (Figure 3I, Table S2), including those previously described (such as *Lyve1* and *Mrc1* in the YS ECs^30^, *Sox9* in trunk ECs of the cardiac cushion^31^, *Hox* genes in the tail ECs, and *Foxl2* in cranial ECs^32^), as well as several genes not previously described as tissue specific in ECs, for example upregulation of *Enpp2* in cranial ECs. We then carried out DGE comparing ECs across different clonal clusters. For this we saw fewer significant DEGs (Table S3), likely due to the low numbers of cells in each embryo. Nevertheless, distinct lineage-associated signatures were detectable. Somitic mesoderm-linked ECs (Mode 5) upregulated *Rbp1* which encodes retinol-binding protein 1, in keeping with the role of retinoid signaling in somitogenesis (Figure 3J). Meanwhile hindlimb-linked ECs (Mode 6) upregulated *Ifitm1* (a marker of posterior mesoderm), and ExM-associated ECs (Mode 3) showed upregulation of *Fmo1*, suggesting distinct metabolic processes in these ECs. The *Rbp1* and *Ifitm1* upregulation was not solely due to anatomical location since their expression was enriched in Modes 5 and 6 even when compared to other modes within the trunk ECs and tail ECs respectively (Figure S12D). Together, these data indicate that while different mesoderm origins can converge to acquire EC identify, there are nevertheless transcriptional differences among ECs that correspond to anatomical position and clonal lineage.

### Allantois has clonal cousins in the extra-embryonic mesoderm and hindlimb

Next, we focused on the lineages giving rise to the extra-embryonic mesoderm (ExM; Figure 4A,B). ExM generates support structures that are essential for embryonic survival: the yolk sac (YS) is the site of early blood formation as well as nutrient and gas exchange; the amnion forms a cushioning layer around the embryo; and the allantois establishes the connection to the placenta and gives rise to the umbilical vessels that link the embryonic and placental circulation. ExM arises from the posterior primitive streak of the early-to mid-streak embryo^3^, within the same territory that generates embryonic mesoderm. However, this shared territory does not establish which extraembryonic and embryonic tissues descend from common progenitors. To investigate this, we first examined the cellular cousins of ExM cell types grouped by mode (as defined in Figure 2) and observed two broad patterns of ExM sharing (Figure 4C, S13A). YS mesenchyme, amniotic mesoderm, and hematopoietic cells shared overlapping clones with one another and with cells in the YS and trunk; we named these cell types “ExM 1”. In contrast, allantois-containing clones were predominantly shared with cells in the YS and tail; we name allantois-containing cells “ExM 2”. Given that 1) YS mesoderm is the earliest cell type emerging from the primitive streak^1^, and 2) cells exit the streak in an anterior-posterior order^1^, we postulate that ExM 1 may represent cells exiting the streak first while ExM 2 may represent cells exiting the streak at a later point in gastrulation.

**Figure 4:**
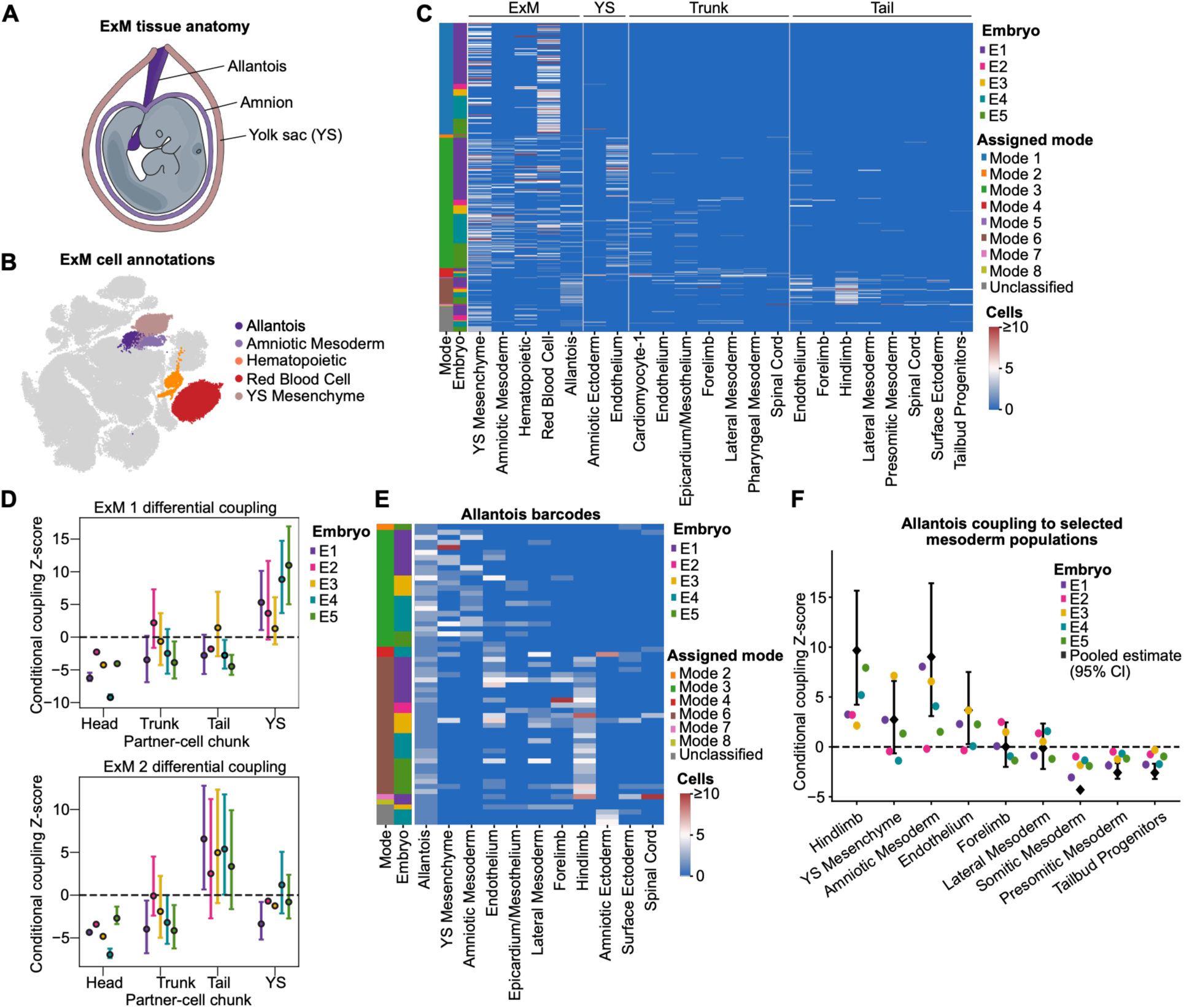
Tracing origins and clonal cousins of extra-embryonic mesoderm lineages. (A) Schematic of extraembryonic mesoderm (ExM) tissues. (B) UMAP showing ExM cell-type annotations. (C) Heatmap showing the distribution of ExM-containing clones across ExM and non-ExM cell-types, grouped by dissection library. Rows represent unique barcodes, ordered by DMM mode assignment and embryo. Color indicates the number of recovered cells, capped at 10. (D) Embryo-specific conditional lineage-coupling Z-scores between ExM 1 (YS mesenchyme, Amniotic Mesoderm, and blood) and 2 (allantois) populations and non-ExM cells from the head, trunk, tail and yolk sac libraries. Points represent individual embryos (E1-E5), with error bars indicating bootstrap 95% confidence intervals. (E) Heatmap showing the cell-type composition of allantois-containing clones grouped by DMM mode assignment. Rows represent individual clones and columns represent cell types; color indicates the number of recovered cells. Annotation bars indicate DMM mode and embryo of origin. (F) Embryo-specific conditional lineage-coupling Z-scores between allantois and selected mesoderm cell types. Points represent individual embryos (E1-E5), with error bars indicating bootstrap 95% confidence intervals. Only clones containing ≥3 recovered cells are included. All panels show data from embryos barcoded at E6.5 (iE6.5).

Taking a closer look at the identity of trunk barcodes found in ExM 1 cells, we observed a surprising enrichment of cardiomyocytes, epicardial cells, and endothelial cells (Figure 4C). These data support the existence of progenitors with dual YS and cardiac fate^33^ and will be discussed in more detail in Figure 5. In contrast, the clonal cousins of ExM 2 cells were diverse: they shared clones with both ExM cells in the YS as well as hindlimb mesoderm in the tail library (Figure 4D,E). To investigate the robustness and specificity of these clonal connections we compared allantois lineage coupling Z-scores across mesodermal cell types across embryos in the iE6.5 dataset. All embryos showed a positive allantois-hindlimb Z-score indicating enriched coupling compared to the null permutations (Figure 4F). Allantois-amniotic mesoderm and allantois-YS mesoderm coupling scores were also largely positive though showed more variation across embryos. Nevertheless, these clonal couplings exceeded that between the allantois and other mesoderm populations, including somitic mesoderm, presomitic mesoderm, and tailbud progenitors which were consistently below the null expectation, while forelimb and lateral mesoderm showed variable coupling across embryos and averaged close to the null expectation. Clone-level bootstrapping revealed substantial uncertainty in some embryo-specific estimates, highlighting variation in the strength of coupling among embryos. Together, these findings identify an enriched clonal relationship between the allantois and hindlimb, as well as allantois and ExM, suggesting the existence of a previously undescribed progenitor that gives rise to the umbilical vessels and embryonic hindlimb mesoderm and convergent origins of the allantois.

**Figure 5:**
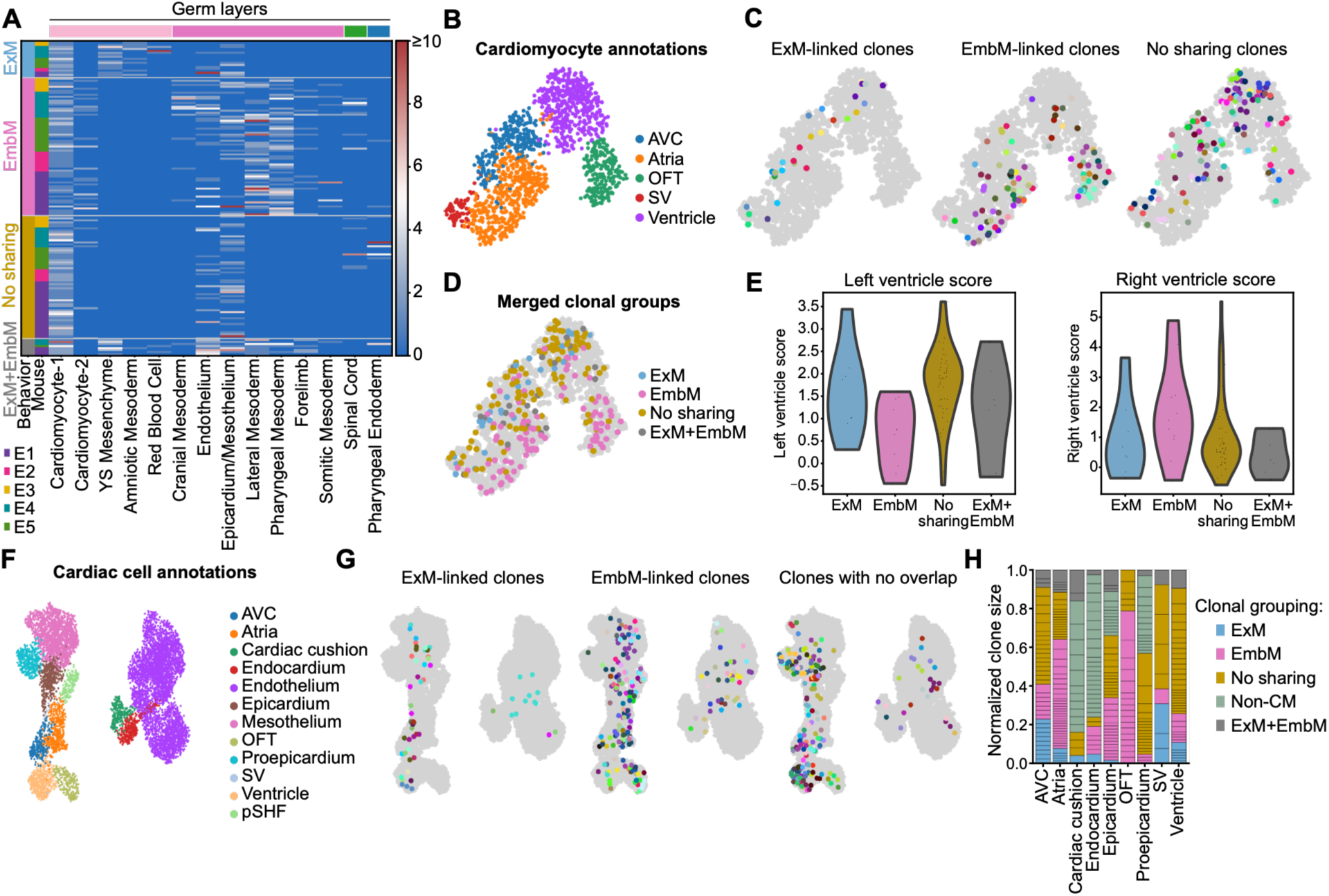
Identifying fate restriction of cardiac progenitors. (A) Heatmap showing the cell-type composition of cardiomyocyte-containing clones, grouped according to clonal sharing with extraembryonic mesoderm (ExM), embryonic mesoderm (EmbM), neither population (No sharing), or both populations (Other) and embryo. Color indicates the number of recovered cells, capped at 10. Only clones containing ≥2 recovered cells are included. (B) UMAP of subclustered and reannotated cardiomyocytes. (C,D) UMAP projections of individual cardiomyocyte-containing clones (C) or clonal sharing categories (D). In C, each color represents a unique barcode. (E) Violin plots showing expression of left-and right-ventricular marker genes across clonal sharing categories in ventricular cardiomyocytes. (F) UMAP of cardiomyocytes, endothelial cells and epicardial cells following subclustering and reannotation. (G) UMAP showing individual clones in different clone-sharing categories, with each color representing a unique barcode. (H) Stacked bar plot showing the distribution of cell types across clonal sharing categories. Only clones containing ≥2 recovered cells are included. AVC, atrioventricular canal; OFT, outflow tract; SV, sinus venosus; pSHF, posterior second heart field. All panels show data from embryos barcoded at E6.5 (iE6.5).

Finally, we identified a small number of ExM clones that contained some cells of the amniotic ectoderm (Figure 4C), that likely arose from a proximal epiblast progenitor^34^. Notably, one of these clones included a primordial germ cell (PGC; Figure S13B,C), and was observed in amniotic ectoderm and allantois lineages. This clone is one of only two multi-cell clones containing PGCs that passed barcode filters across our datasets. The other was observed in the iE5.5 dataset and was shared with hematopoietic cells (myeloid and red blood cells). Although more data will be required to determine whether these associations are reproducible, these data raise the possibility of a shared epiblast origin for extra-embryonic mesoderm, extra-embryonic ectoderm, blood, and PGCs. Interestingly, this clonal relationship has also been reported in a recent preprint that uses a distinct barcoding approach^18^.

### Mapping the distinct clonal origins of cardiomyocytes

Finally, we investigated the clonal origins of cardiomyocytes (CMs). Existing models propose that CMs arise from at least three different progenitor populations: the first heart field (FHF), second heart field (SHF), and more recently described juxta-cardiac field (JCF)^35,36^. These populations make distinct contributions to different chambers and structures of the developing heart, but questions remain regarding their precise fate boundaries and clonal relationships with other embryonic lineages during gastrulation. In particular, the extent to which cardiac progenitors are restricted to cardiomyocyte fates or share common ancestry with non-cardiac lineages, including endothelial and epicardial cells, remains incompletely understood. As described above, our analysis of ExM lineages highlighted clonal sharing with cardiomyocytes, suggesting a close clonal relationship between the ExM and a subset of cardiomyocytes, as recently reported^33^. Supporting this, analysis of DMM mode assignment demonstrated that although the majority of cardiomyocyte clones are assigned to Mode 4, which is additionally composed of pharyngeal, lateral and forelimb mesoderm, a subset of cardiomyocytes clones are assigned to Mode 3, which is enriched in ExM (Figure 2D). Furthermore, all embryos in the iE6.5 dataset contained cardiomyocytes that were assigned to both Mode 3 and 4 (Figure S14A). Since DMM modes capture broad patterns of clonal composition rather than the presence or absence of specific lineage relationships, we next directly examined the clonal cousins of cardiomyocytes. Specifically, we classified CM-containing clones according to whether they shared barcodes with ExM (YS mesenchyme, amniotic mesoderm, and blood) or EmbM (pharyngeal, lateral, cranial, limb, and somitic mesoderm) populations. This revealed that ∼11% of CMs shared barcodes with ExM (18/157 CM-containing clones), while ∼39% shared barcodes with EmbM (69/157 clones; Figure 5A). A further 42% of CMs showed no detectable sharing with either group, while 8% shared barcodes with both (Figure 5A). Given that clonal sharing with ExM and EmbM was largely non-overlapping among recovered clones, we hypothesized that these patterns might reflect distinct progenitor routes contributing to cardiomyocytes.

To determine whether ExM, EmbM, no-sharing, or both groups were observed in different cardiac regions, we annotated CMs into ventricular, atrial, outflow tract, and atrioventricular canal (AVC) identities using established markers (Figure 5B, S14B)^37^. Projection of ExM, EmbM and no-sharing clones revealed distinct spatial segregation: ExM-shared clones contributed primarily to the ventricle and AVC, whereas EmbM-shared clones were enriched in the outflow tract, atria, and ventricle (Figure 5C,D, S14C). No-sharing clones overlapped most closely with ExM-shared cells and were abundant in the ventricle. Given that the left and right ventricle are derived from different lineages (FHF and SHF, respectively^32,33^), we investigated whether the clonal groups were associated with ventricular CMs with distinct transcriptional signatures. This revealed that ventricular CMs associated with ExM and no-sharing clones express enriched levels of previously described left ventricle markers^37^, while ventricular CMs associated with EmbM express enriched levels of previously described right ventricle markers^37^ (Figure 5E). These findings indicate that analysis of CM clonal cousins can reveal distinct fate behaviors that resemble previously described JCF/FHF (ExM and no-sharing groups) and SHF (EmbM group) lineages.

We finally examined broader progenitor fate by analyzing clone sharing between CM and non-CM cardiac populations. For this we combined all cells labelled as “endothelial” or “epicardial” in our original dataset in addition to CMs. Subclustering enabled us to further identify epicardium, proepicardium, endocardium, cardiac cushion, and mesothelium cells using previously described markers (Figure 5F, S14B)^37,38^. Notably, CM-associated cells represented a minority of cells within the endocardium (<25% cells; Figure 5G,H, S2A, S14D). In contrast, 60-77% proepicardium and epicardial cells belonged to clones that were shared with CMs, supporting a stronger clonal connection between CMs and epicardium than CMs and endocardium at the time of labelling (Figure 5H), in keeping with recent work^33,39,40^. Together, these findings reveal distinct clonal routes to cardiomyocyte formation that are associated with specific cardiac identities, and demonstrate that early progenitors contributing to cardiomyocytes can also give rise to other cardiac lineages, particularly the epicardium.

## Discussion

Here, we use CRISPR-based barcoding to perform high-resolution lineage tracing of the early mammalian embryo at the onset of gastrulation, enabling us to quantify lineage relationships across germ layers and map clonal diversification at organismal scale. A unifying theme emerging from our analysis is that spatially and functionally distinct progenitors converge to generate transcriptionally similar cell types. We observe this across multiple mesodermal lineages, including the somitic mesoderm (Figure 2), endothelium (Figure 3), allantois (Figure 4), and cardiomyocytes (Figure 5). This finding has important implications across developmental and regenerative biology, including for understanding the origins of cell heterogeneity within tissues and informing the design of *in vitro* organogenesis strategies, which may benefit from capturing heterogeneity of cell origins. Furthermore, while our focus here is on the mesoderm, our dataset contains ectodermal and endodermal cells with robust barcode labelling (Figures S2,4), and can be explored through our datasets and interactive website.

Our data provide direct clonal evidence supporting a dual origin of paraxial mesoderm, in keeping with previous studies based on clonal tracing, fate mapping, and lineage inference^27^. The anterior enrichment of mesoderm-biased clones within the paraxial mesoderm supports the existence of two sources of somitic mesoderm: one directly from the primitive streak and another from tailbud-resident NMPs that have been proposed to generate neural and paraxial mesoderm along the elongating embryonic axis^6,26^. However, given our early labelling timepoint, we cannot exclude the possibility that some clones containing both mesoderm and ectoderm cells represent epiblast cells labelled prior to fate restriction as previously described^5^. Regardless, these findings support a model in which transcriptionally similar somitic mesoderm cells arise from multiple developmentally distinct progenitor pools, highlighting the convergence of lineage trajectories during mammalian axis formation.

A similar principle is evident in endothelial development, where we observe convergence of a shared transcriptional state from multiple mesodermal progenitors. Our barcoding studies confirm that ECs can arise from multiple different types of mesoderm, building on classical studies demonstrating that ECs separately emerge from lateral, extra-embryonic, and paraxial mesoderm^41–44^. Notably, these diverse origins have not previously been predicted through trajectory analyses of transcriptional data^10,32^, highlighting the importance of ground-truth lineage tracing data for establishing clonal ancestry. In addition to previously described regional heterogeneity^32^, we show that transcriptional variation among ECs is linked to their clonal origins, suggesting that early lineage segregation may contribute to tissue-specific endothelial diversity (Figure 3J). More broadly, this raises the possibility that developmental origin is a driver of cellular heterogeneity across embryonic and adult tissues.

Our data also indicate that the allantois, the predecessor of the umbilical cord that connects the placenta and embryo, is generated from at least two distinct progenitor pools: one associated with extra-embryonic mesoderm, and another associated with the hindlimb. Here, our data supports previous fate-mapping studies that have demonstrated that the allantois arises from progenitors associated with both embryonic and extra-embryonic mesoderm^3^, though the specific embryonic lineages associated with allantois formation have not previously been characterized. Our identification of preferential allantois-hindlimb clonal sharing raises the possibility that the allantois receives contributions from two regions of the primitive streak, potentially reflecting distinct phases of allantois formation. Interestingly, the hindlimb and allantois share expression of Tbx4 (Figure S2) which has previously been recognized but not associated with a common developmental origin^45^.

Finally, clonal analysis of cardiomyocytes reveals contributions from multiple progenitor populations to distinct cardiac subtypes, in keeping with current models of cardiac development^35^, Based on our observed lineage patterns, we propose that EmbM-linked clones correspond to SHF progenitors given their lineage overlap with pharyngeal, cranial, and lateral mesoderm. Meanwhile, we propose that ExM-linked clones correspond to recently described JCF-like progenitors given their fate outcomes (contribution to left ventricular CMs, AVC myocardium, and epicardium) and sharing with YS mesoderm^33,40^. While the no-sharing lineage is likely to encompass cell types that do not meet the EmbM/ExM sharing criteria as a result of incomplete barcode sampling, another possibility is that it is enriched with FHF-like progenitors, since these cell types were associated with LV and AVC CM fates, and FHF progenitor fate has previously been reported to diverge early from other mesoderm cells during gastrulation^46^. Together, these findings demonstrate how combined clonal and transcriptional approaches can resolve progenitor contributions to cardiac cell diversity.

### Limitations of the study

Our dataset is limited by incomplete editing and sampling of barcodes and cells. From each mouse, we sampled ∼4-8% of the mouse embryo for scRNA-seq analysis based on previous cell number estimates^9^. Given a useable barcode was obtained from an average of ∼15% cells across all datasets, approximately 0.2-1.2% of embryonic cells were sampled (breakdown shown in Figure S3G). Future work would benefit from use of the DARLIN model (which in our dataset results in higher editing efficiency averaging 56% compared with CARLIN embryos that averaged 37%), use of three barcode arrays (here we only use barcodes from the *Col1a1* and *Tigre* loci and did not analyze *Rosa26* locus barcodes^20^), and from increased sampling of the mouse embryo through sequencing of more cells. Finally, the datasets generated here represent fate maps and therefore provide no indication of cell potential at the time of labelling. Tissue grafting experiments have demonstrated that the potential of embryonic cells are greater than their fate at the onset of gastrulation^47,48^, highlighting that during gastrulation cell fate allocation is not concomitant with fate determination.

## Acknowledgements

The authors wish to thank members of the Bowling, Camargo, and Göttgens laboratory for discussion. We are grateful for funding from: National Institutes of Health grant T32GM141828 (PJM, PJD); Schmidt Science Fellows (JCS); Isaac Newton Trust Junior Research Fellowship (JCS); Wolfson College Junior Research Fellowship (JCS); National Institutes of Health grant 1K99HL164969 (SJB); Wellcome Trust grant 215920/Z/19/Z (SJB); European Molecular Biology Organization Fellowship ALTF 798-2018 (SJB); National Institutes of Health grant R01HL128850 (FDC); National Institutes of Health grant RC2DK131963 (FDC); National Institutes of Health grant R01HL158192 (FDC); Edward P Evans Foundation (FDC); Alex Lemonade Crazy 8 award (FDC).

## Author Contributions

Conceptualization: BG, FDC, SJB; Experiments: MAS, PJM, SJB; Analysis: MAS, PJM, JCS, DVE, PJD, EYL, NL, SJB; Statistical modelling: JCS; Annotations: MAS, PJM, AM, HDK, RTK, KML, SJB; Funding acquisition: FDC, SJB; Supervision: XQ, KML, BG, FDC, SJB; Writing – original draft: MS, JCS, SJB; Writing – review & editing: MAS, JCS, PJM, KML, BG, FDC, SJB.

---

**Figure S1:**
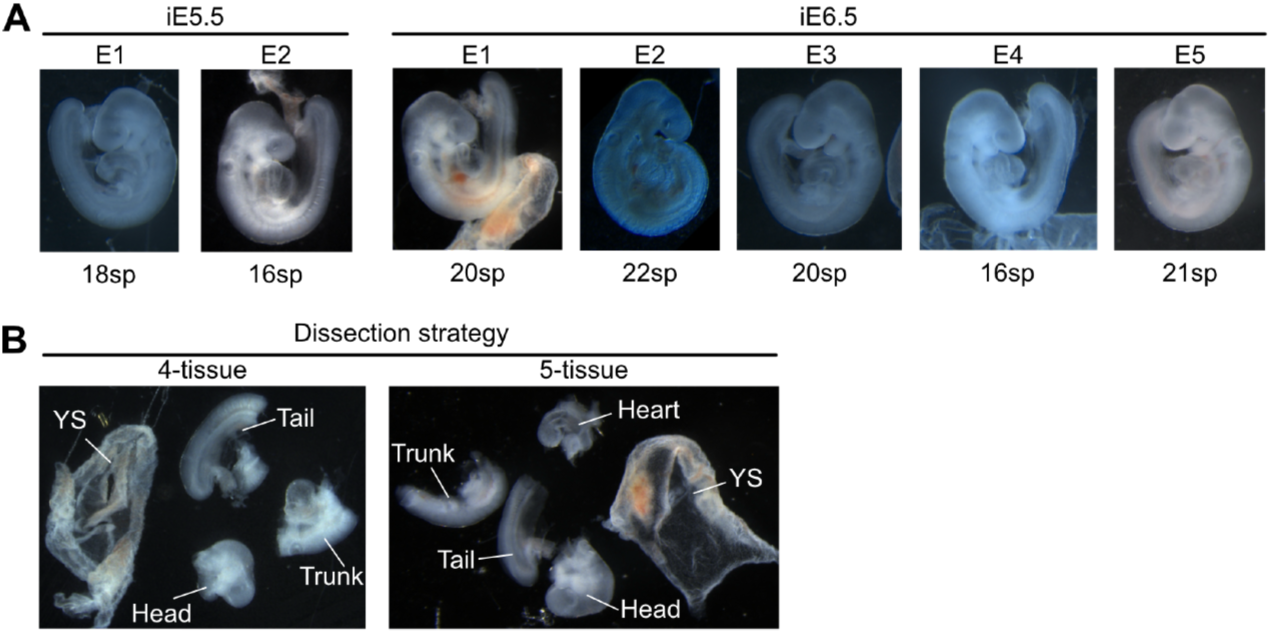
Barcoded embryos used for scRNA-seq analysis. (A) Images of all E9.5 embryos collected for scRNA-seq analysis; corresponding somite numbers (sp) are indicated below each embryo. (B) Dissection strategy used for the embryos. All embryos were dissected into four regions (left), apart from E1 which was dissected into five regions (right).

**Figure S2:**
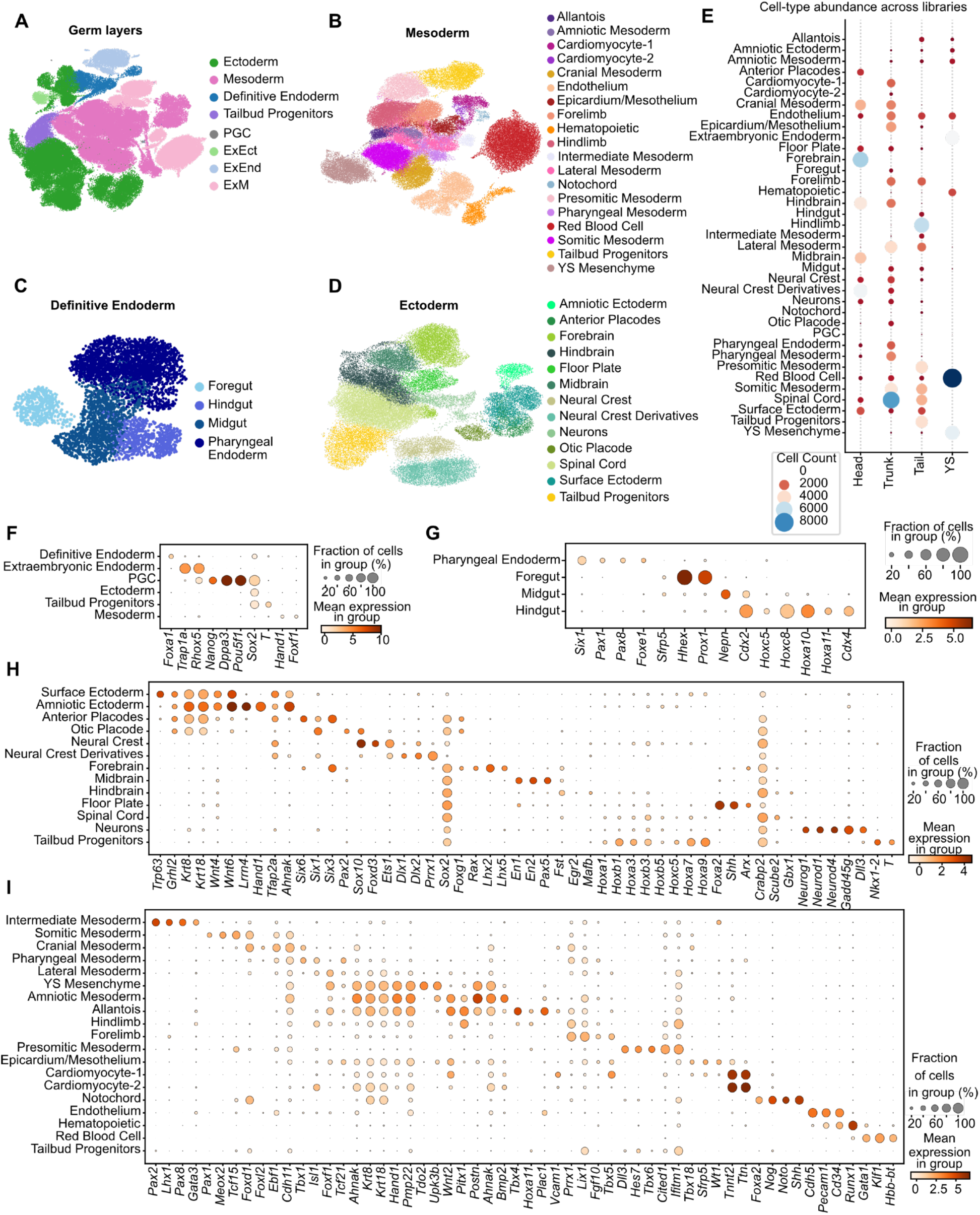
Cell type annotations. (A-D) UMAP colored by the germ layer annotations (A), mesoderm cell types (B), definitive endoderm cell types (C), ectoderm cell types (D). E. Distribution of cell types across the different tissue libraries. (F-I) Dot plots showing expression of marker genes used to annotate germ layers (F), endoderm (G), ectoderm (H) and mesoderm (I).

**Figure S3:**
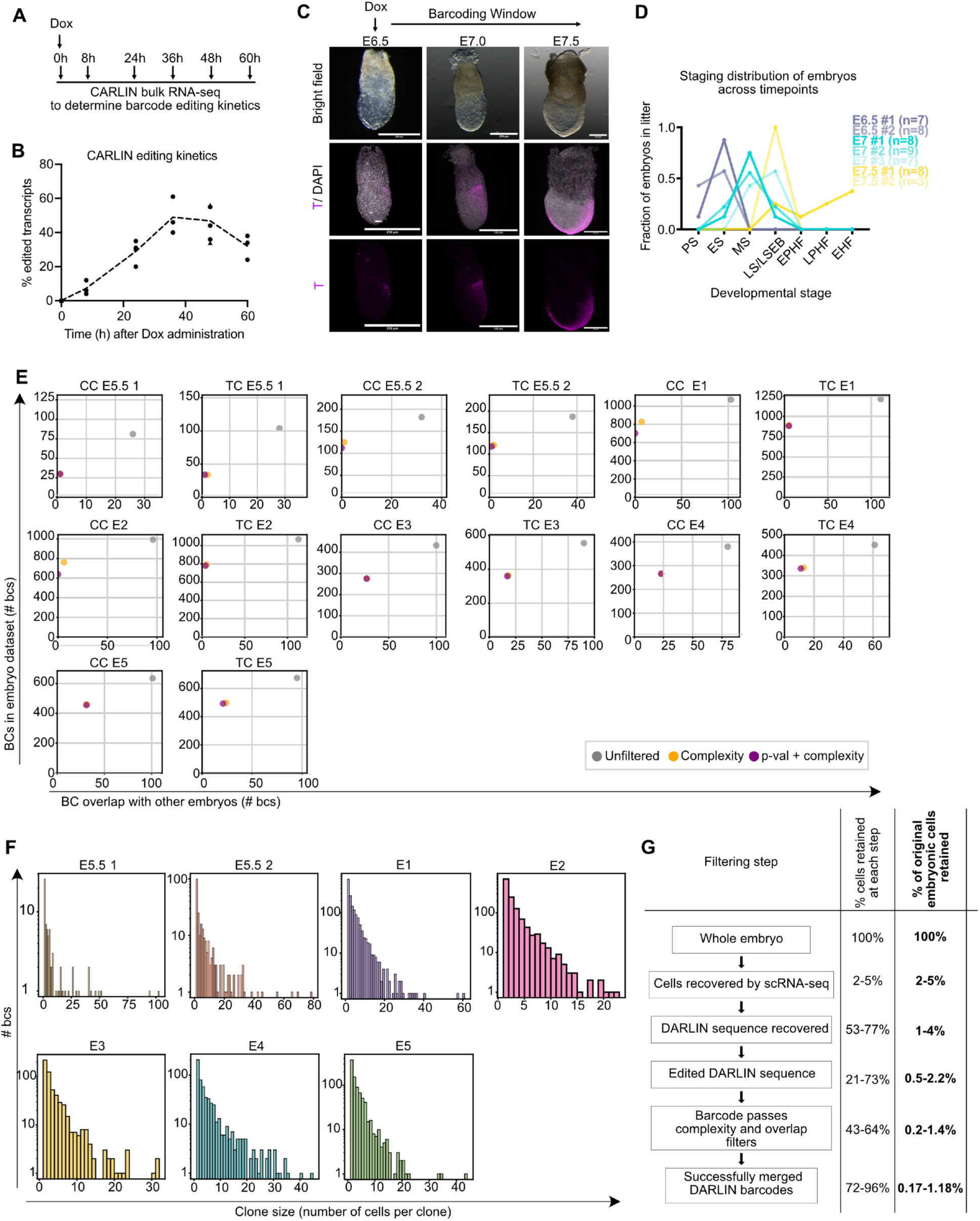
Barcode editing kinetics, filtering and clone-size distributions. (A) Schematic of experimental design for quantification of CARLIN barcode editing kinetics. Doxycycline (Dox) was administered at 0 h, and embryos were collected at 8, 24, 36, 48 and 60 h for bulk RNA sequencing of CARLIN transcripts. Uninduced control embryos (plotted as 0 h) were collected at E8.5 and E9.5. Sample sizes were: 0 h, 2 litters/6 embryos; 8 h, 1 litter/3 embryos; 24 h, 2 litters/4 embryos; 36 h, 1 litter/3 embryos; 48 h, 2 litters/6 embryos; and 60 h, 1 litter/3 embryos. (B) Percentage of CARLIN transcripts containing edited barcodes at each time point following Dox administration. Points represent individual embryos; the dashed line indicates the trend in editing efficiency over time. (C) Representative images of uninduced DARLIN embryos collected at E6.5, E7 and E7.5 and stained for Brachyury (T) to assess developmental stage. (D) Proportion of embryos corresponding to different developmental stages within each litter. Each line represents an individual litter, with the number of embryos indicated. PS, primitive streak; ES, early streak; MS, midstreak; LS, late streak; LSEB, late streak/early bud; EPHF, early pre-headfold; LPHF, late pre-headfold; EHF, early headfold. (E) Barcode overlap between embryos at the Col1a1 (CC) and Tigre (TC) barcode arrays under different filtering strategies. The complexity filter was used for all subsequent analyses. (F) Distribution of recovered clone sizes across individual embryos. (G) DARLIN barcode recovery across sequential filtering steps. Percentages indicate the minimum-maximum range across embryos; individual embryo statistics are provided in Table S1. Whole-embryo cell numbers were estimated rather than directly measured.

**Figure S4:**
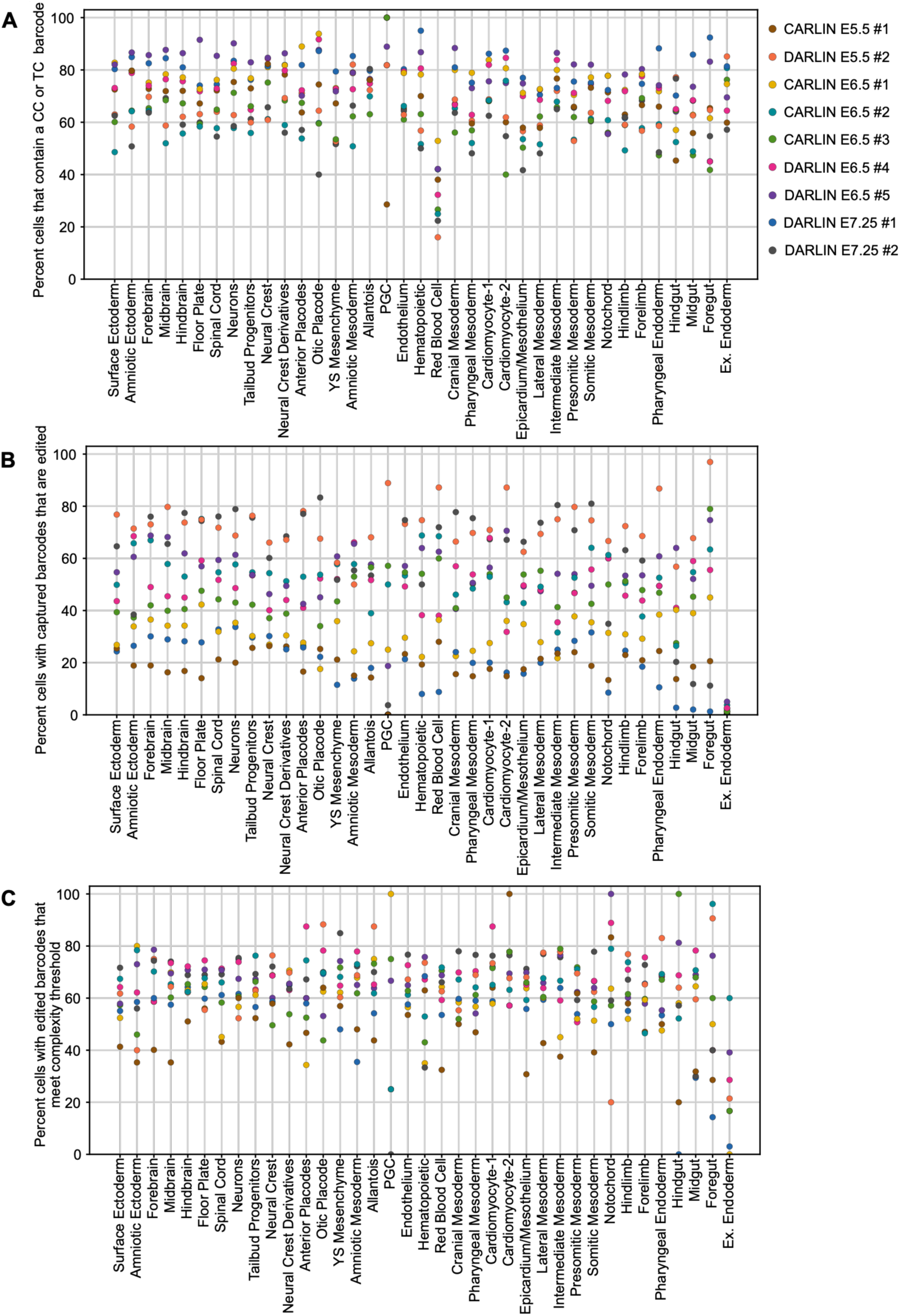
Barcoding efficiency and average clone size per cell type. (A) Barcode capture efficiency across cell types in individual embryos. (B) Percentage of cells with captured barcodes containing edits, grouped by cell type and embryo. (C) Percentage of cells with edited barcodes passing the complexity filter, grouped by cell type and embryo.

**Figures S5:**
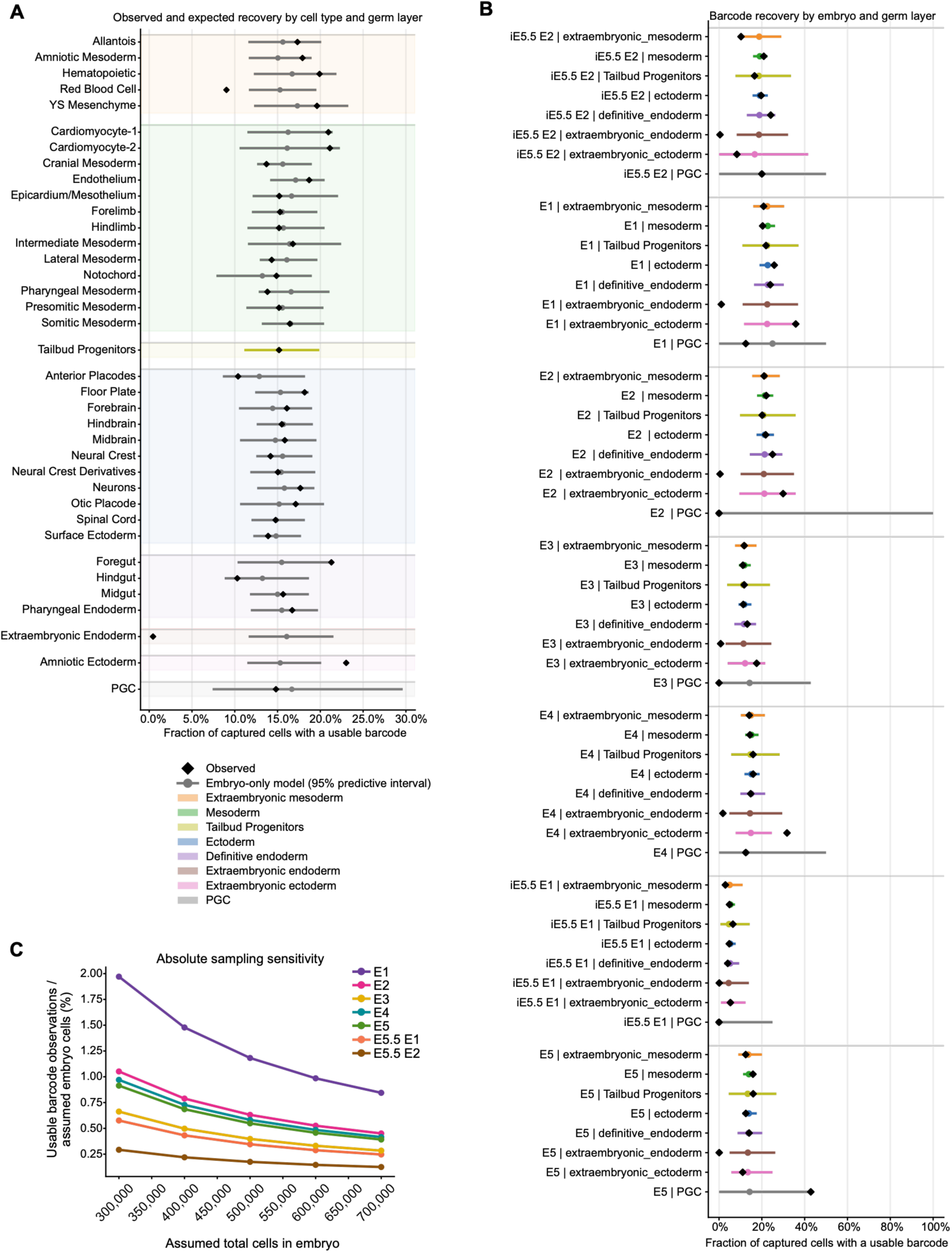
Observed barcode recovery across cell types and embryos compared with an embryo-specific sampling model. (A) Observed proportion of captured cells with usable DARLIN barcodes across cell types, pooled across embryos (black diamonds), compared with predictions from an embryo-specific beta-binomial model. Grey circles indicate model expectations and horizontal lines indicate 95% predictive intervals. (B) Observed barcode recovery (black diamonds) and model predictions (colored circles) across germ layers and developmental populations within individual embryos. Horizontal lines indicate 95% predictive intervals. The model accounts for embryo-specific variation in barcode recovery and overdispersion, providing a reference for assessing cell-type- and lineage-specific deviations. (C) Modelled usable barcode recovery proportion across total embryo cell counts per embryo.

**Figure S6:**
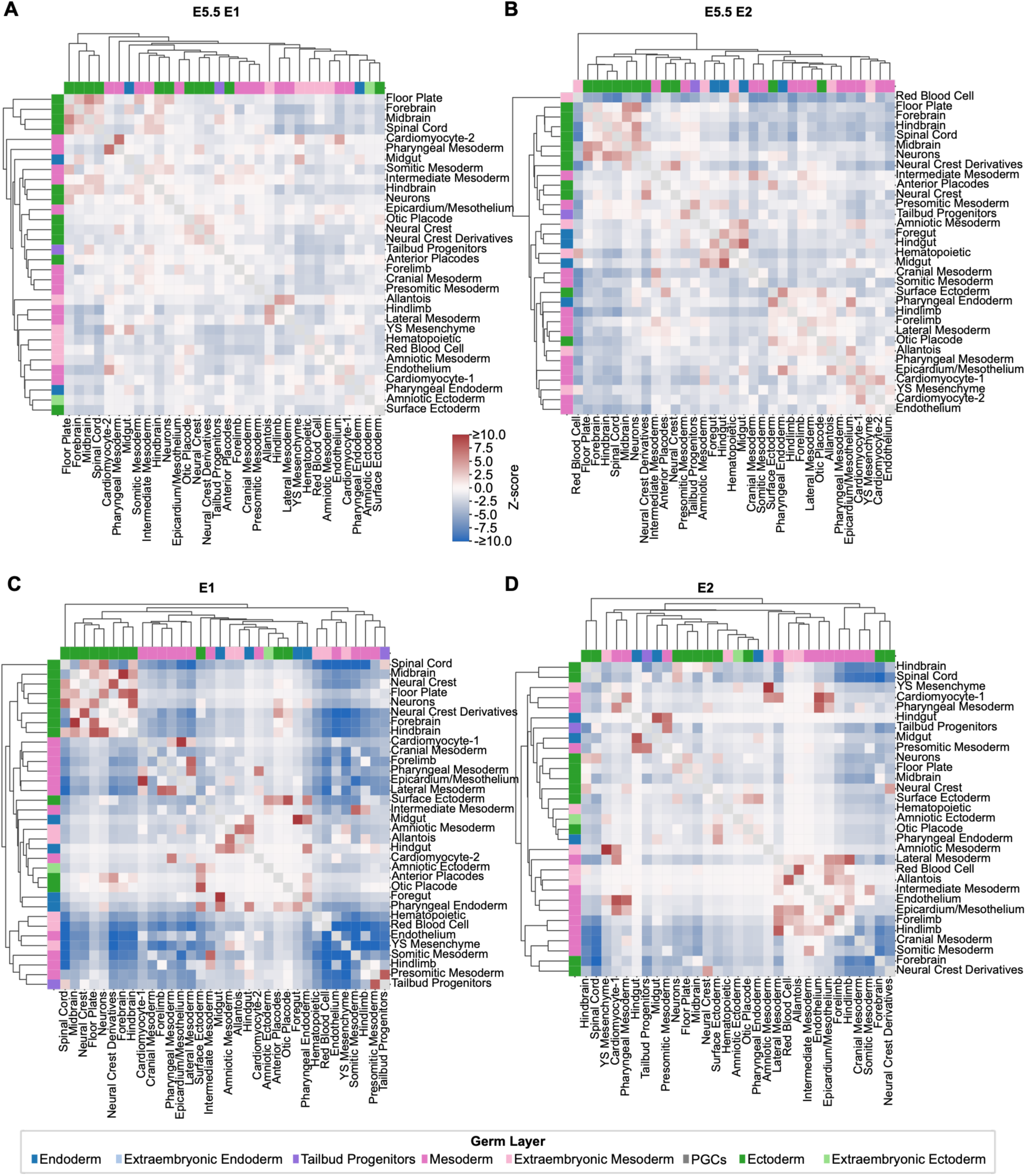
Lineage coupling scores for individual embryos. (A-D) Heatmaps showing lineage-coupling Z-scores across cell types in individual embryos induced at E5.5 (A,B) or E6.5 (C,D; E1 and E2). Z-scores were calculated relative to null distributions generated from 10,000 permutations. Cell types are colored according to germ-layer annotation, using the key shown in Figure 1. Only clones containing ≥3 recovered cells are included.

**Figure S7:**
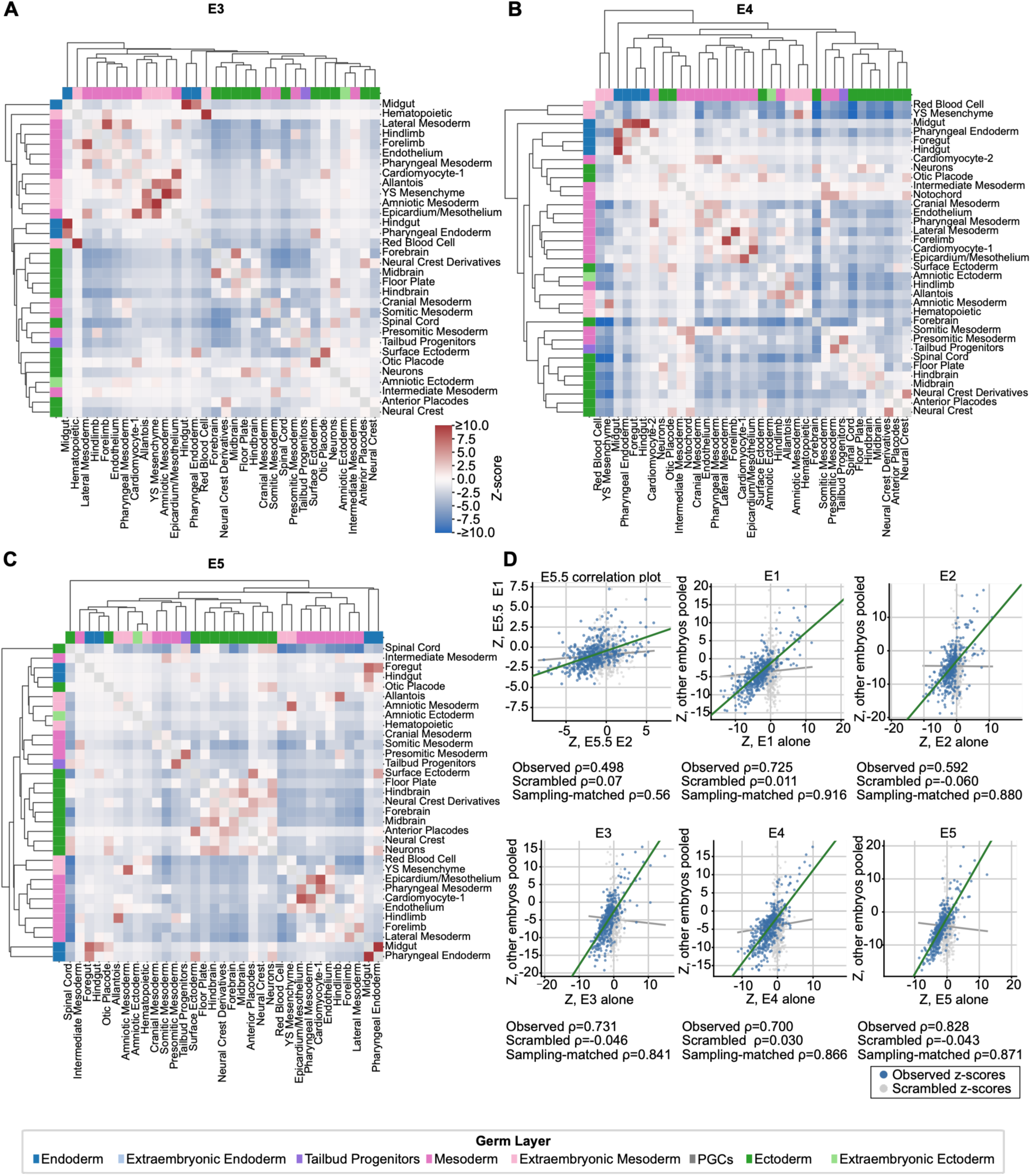
Lineage coupling scores and reproducibility across embryos. (A-C) Heatmaps showing lineage-coupling Z-scores across cell types in individual embryos induced at E6.5 (E3-E5). Z-scores were calculated relative to null distributions generated from 10,000 permutations. Cell types are colored according to germ-layer annotation, using the key shown in Figure 1. Only clones containing ≥3 recovered cells are included. (D) Spearman correlations between lineage-coupling Z-scores calculated for individual iE6.5 embryos and the pooled remaining embryos (observed ρ, blue), compared with scrambled (grey) controls. Scrambled controls were generated by permuting cell-type labels within individual embryos. Sampling-matched controls were generated by drawing two disjoint sets of clones from the pooled dataset, matched in size to the individual embryo and the remaining embryos, respectively. For the subsampled controls, the median correlation from 10 iterations is shown. Each point represents a cell-type pair; lines indicate least-squares regression fits. Correlations were calculated using 465 cell-type pairs represented in all embryos.

**Figure S8:**
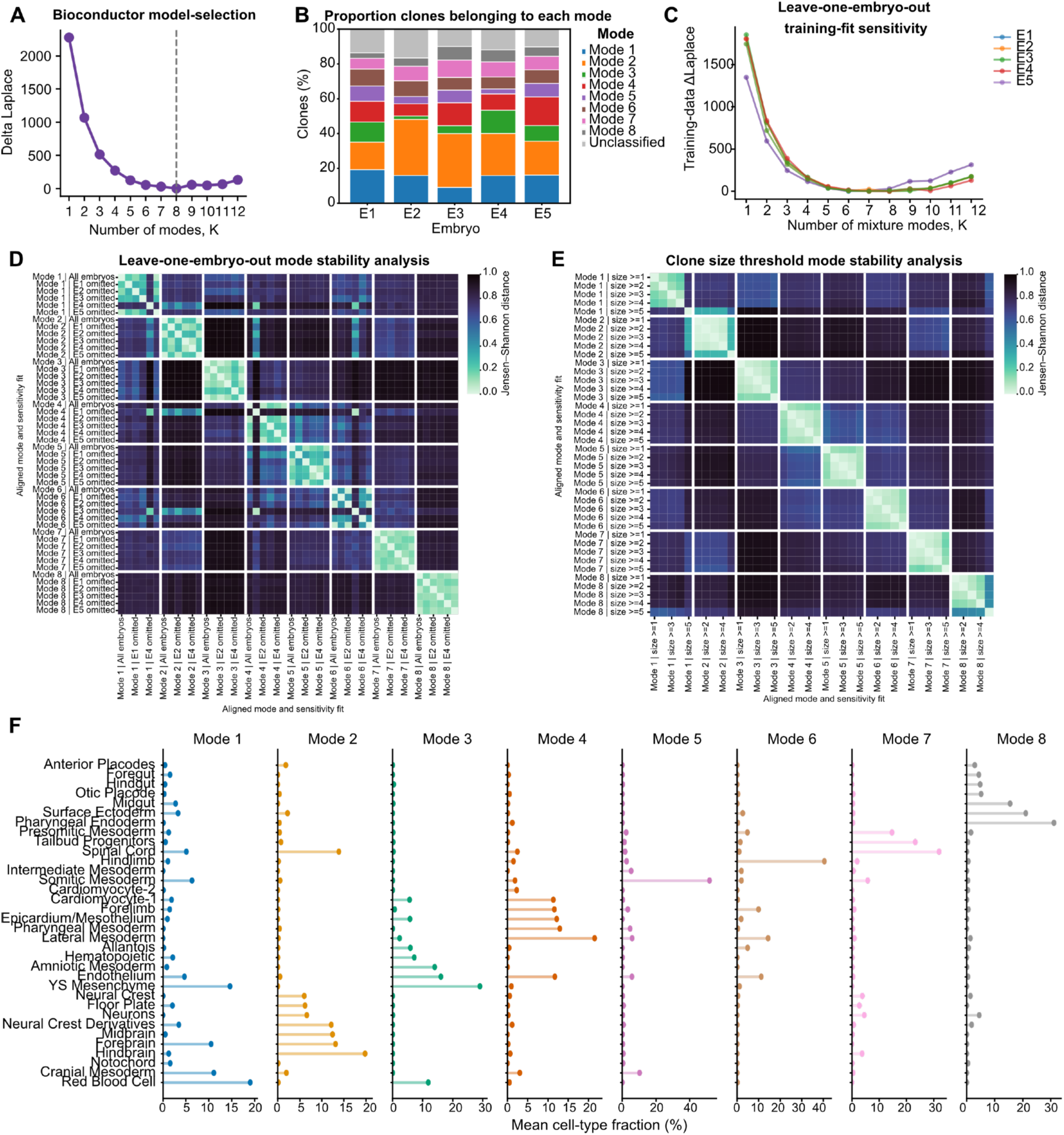
Dirichlet-Multinomial model selection and robustness. (A) Model selection across different numbers of Dirichlet-multinomial mixture (DMM) modes (K = 1-12), based on the Laplace approximation to model evidence. The dashed line indicates the selected eight-mode model. (B) Distribution of DMM mode assignments across individual embryos. Stacked bars indicate the proportion of clones assigned to each of the eight modes or classified as unclassified. Clones were assigned to a mode if its posterior probability was at least twice that of the next most probable mode. Only clones containing ≥3 recovered cells are included. (C) Leave-one-embryo-out analysis showing the sensitivity of model selection to the exclusion of individual embryos. Lines indicate the training-data Laplace criterion across different numbers of modes, with each embryo excluded in turn. (D) Leave-one-embryo-out mode stability analysis. Heatmap showing pairwise Jensen-Shannon distances between mode contribution profiles inferred from the pooled dataset and those inferred with individual embryos excluded. (E) Clone-size threshold sensitivity analysis showing pairwise Jensen-Shannon distances between mode contribution profiles inferred using different minimum recovered clone-size thresholds (≥1-5 cells). (F) Model-estimated cell-type contribution profiles for each of the eight DMM modes. All panels show data from embryos barcoded at E6.5 (iE6.5).

**Figure S9:**
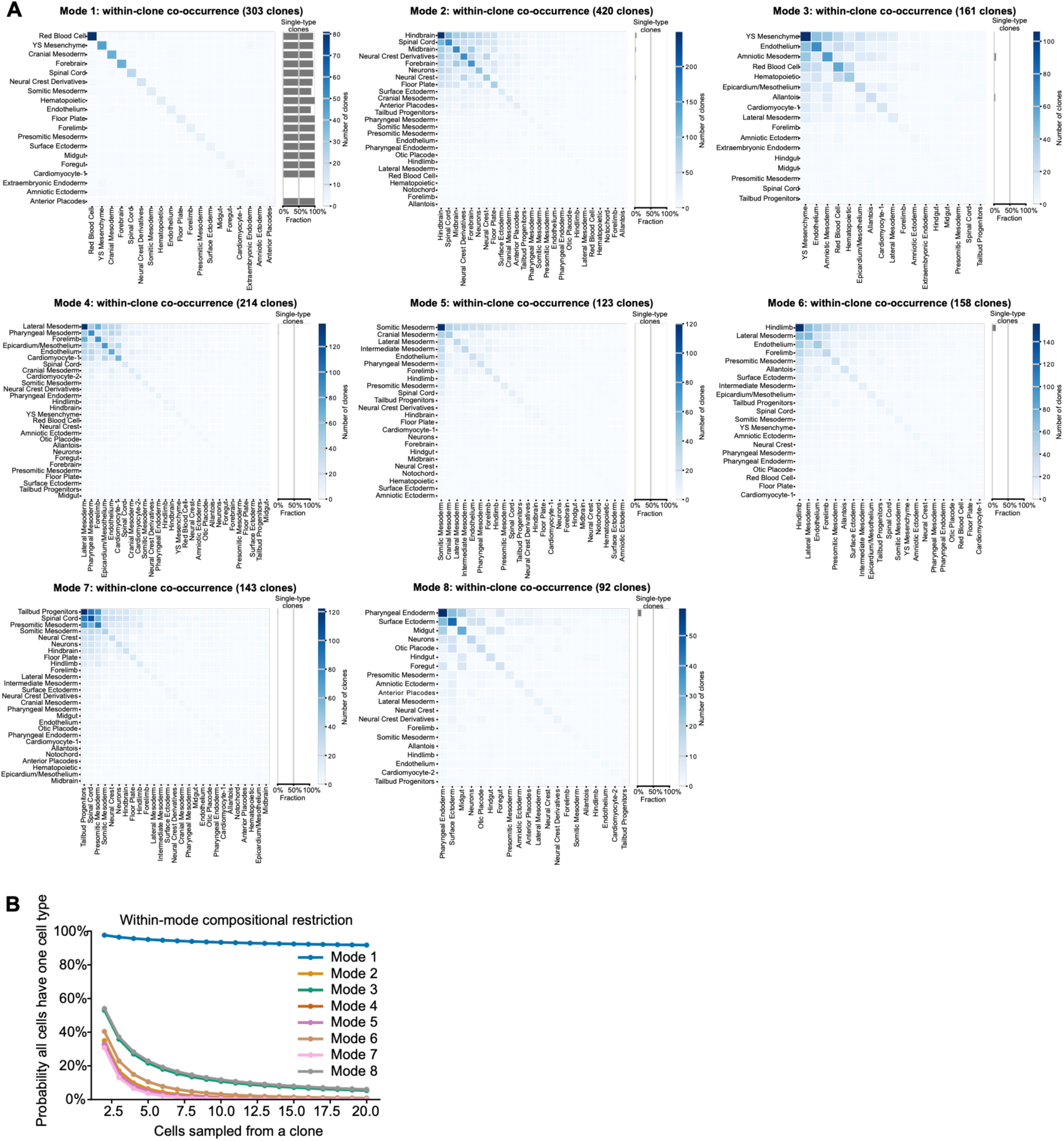
Composition analysis of DMM-derived modes. (A) Heatmaps showing clonal sharing between cell types within each of the eight DMM modes. Diagonal values indicate the number of clones containing each cell type, while off-diagonal values indicate the number of clones containing both cell types. Cell types are ordered according to their frequency within each mode. Adjacent bars indicate the proportion of clones containing only a single cell type. (B) DMM model estimations of the probability that all recovered cells within a clone belong to the same cell type, plotted against the number of recovered cells per clone, grouped by DMM mode. All panels show data from embryos barcoded at E6.5 (iE6.5).

**Figure S10:**
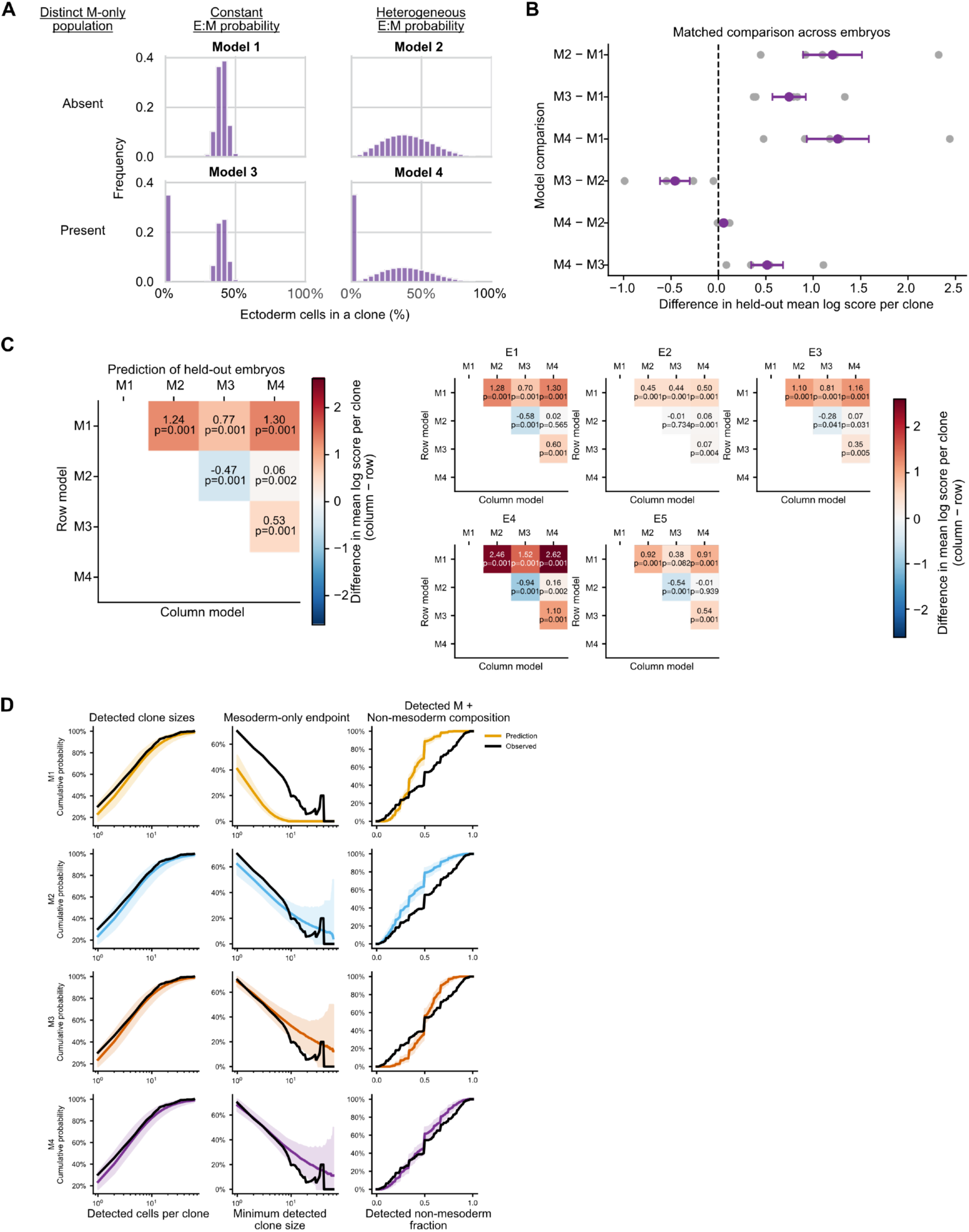
Sampling-aware modelling of clonal contributions to the paraxial mesoderm. (A) Schematic distributions of underlying clone compositions assumed by four generative models (M1-M4). (B) Pairwise comparison of model predictive performance using leave-one-embryo-out cross-validation. Values indicate differences in mean held-out log predictive score per clone, with positive values favoring the first model in each comparison. Grey points represent individual embryos; purple points and intervals indicate pooled estimates and 95% bootstrap confidence intervals. (C,D) Pairwise comparisons of held-out model predictive performance, pooled across embryos (left) and shown separately for each embryo (right). (D) Posterior predictive checks comparing observed data (black lines) with predictions from the four generative models (M1-M4; colored lines and shaded intervals). Models were evaluated based on the distribution of detected clone sizes (left), the probability of detecting no non-mesodermal cells as a function of detected clone size (middle), and the distribution of non-mesodermal fractions among detected mixed clones (right). Colored lines indicate median model predictions, and shaded regions indicate 95% predictive intervals. All panels show data from embryos barcoded at E6.5 (iE6.5).

**Figure S11:**
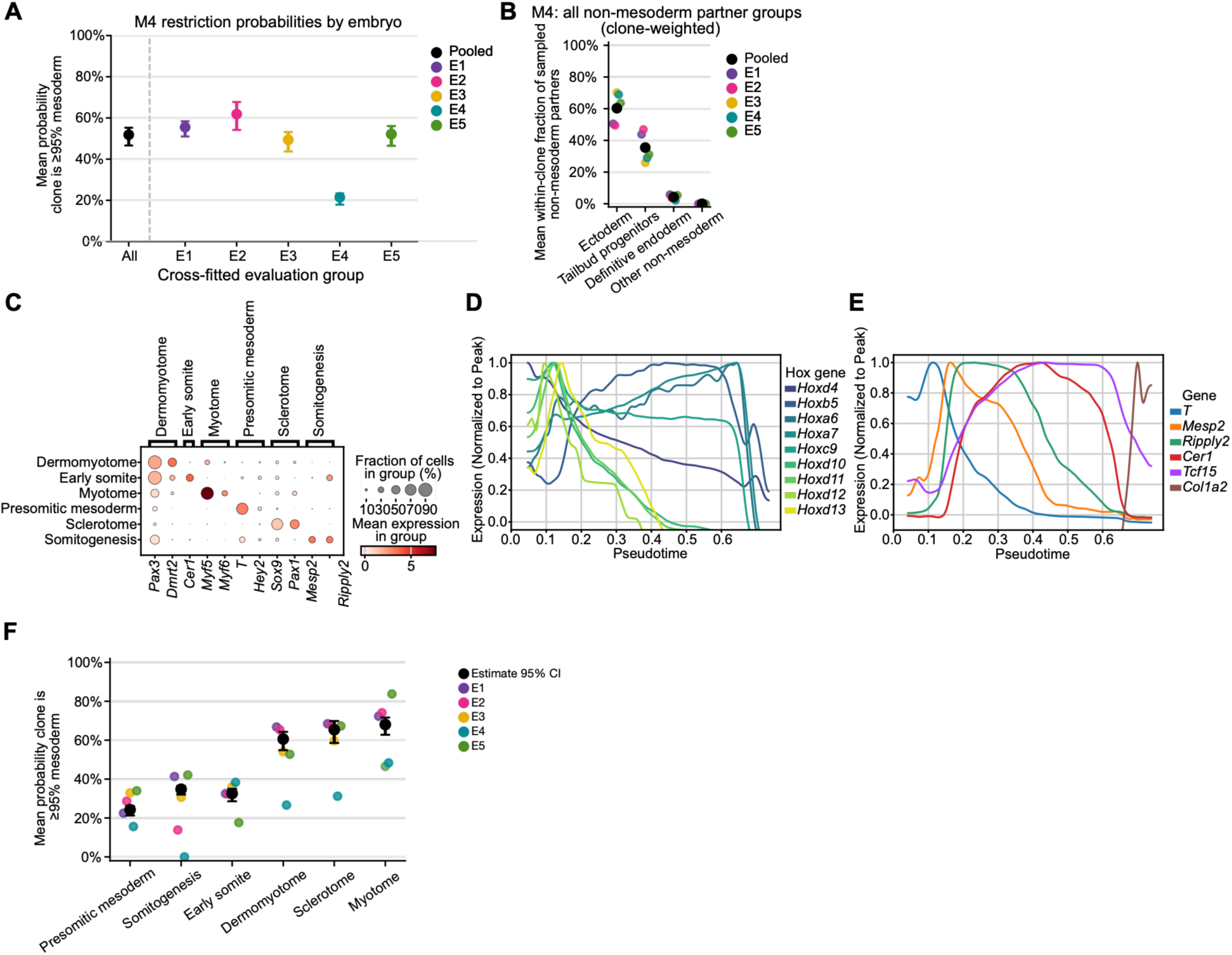
Supplementary analysis of somitic mesoderm origins. (A) Model-estimated probability that sampled mesoderm cells belong to strongly mesoderm-biased clones (≥95% mesoderm; hereafter Mes-biased), using the sampling-aware generative model M4. Points represent individual embryos. (B) Estimated probability of detecting clonal sharing with different non-mesodermal populations among clones inferred to contain less than 95% mesoderm, calculated per clone. Points represent individual embryos. (C) Dot plot showing expression of marker genes used to annotate presomitic and somitic mesoderm populations. (D,E) Normalized expression of Hox genes (D) and somitogenesis-associated genes (E) across diffusion pseudotime. (F) Model-estimated probability that sampled mesoderm cells from individual paraxial mesoderm subtypes belong to strongly mesoderm-biased clones (≥95% mesoderm). Cell types are arranged approximately along the anterior-posterior developmental trajectory, from presomitic mesoderm to differentiated somitic derivatives. Points represent individual embryos. All panels show data from embryos barcoded at E6.5 (iE6.5).

**Figure S12:**
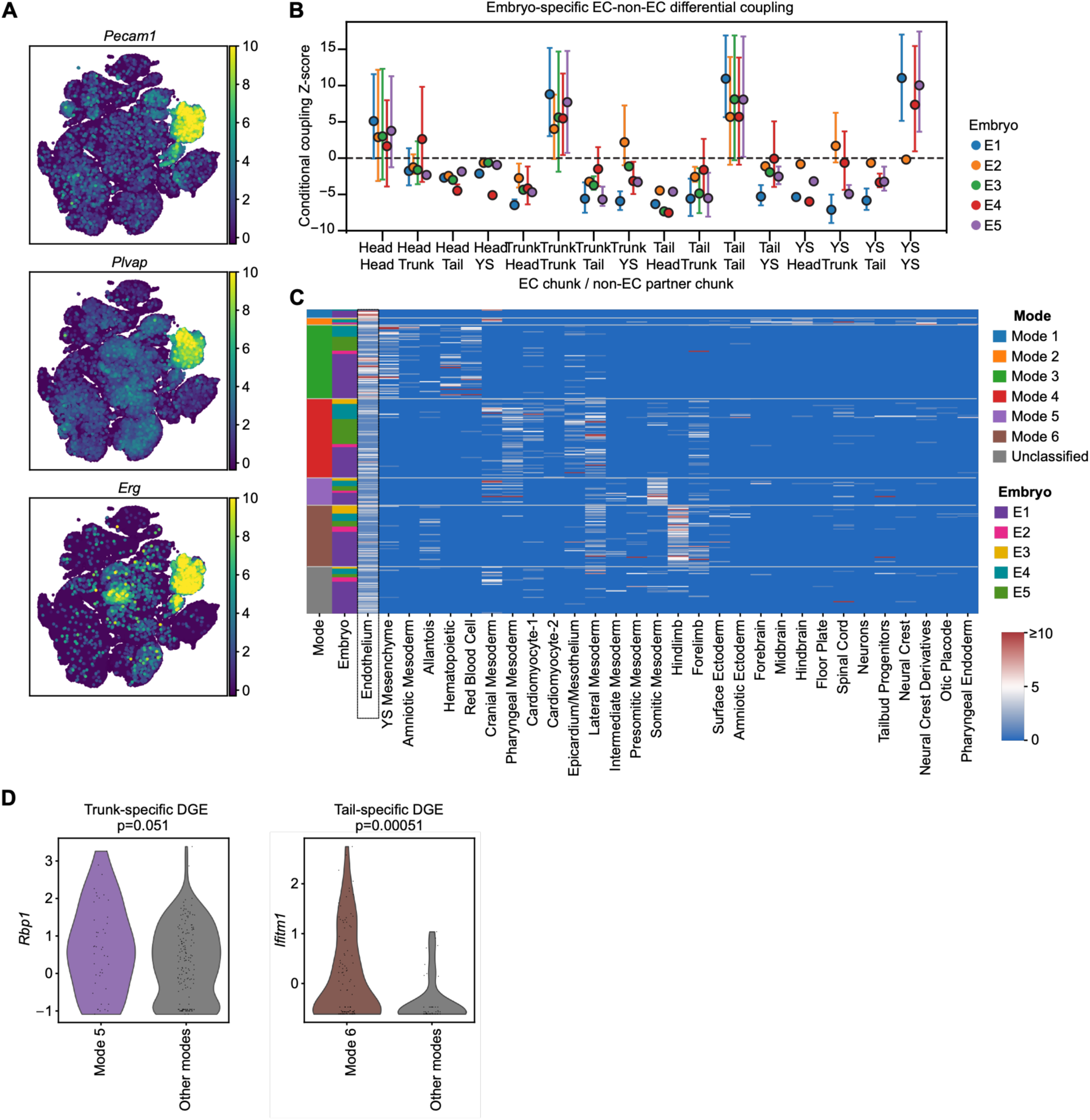
Supplemental analysis of endothelium origins. (A) UMAP showing expression of *Pecam1*, *Plvap* and *Erg* across the transcriptomic dataset. (B) Embryo-specific conditional lineage-coupling Z-scores between ECs and non-EC populations from the head, trunk, tail and yolk sac (YS) libraries. Error bars indicate bootstrap 95% confidence intervals estimated by resampling clones. (C) Heatmap showing the cell-type composition of endothelial cell (EC)-containing clones across cell types. Rows represent unique barcodes, ordered by assigned mode, and columns represent cell types. Color indicates the number of recovered cells, capped at 10. Only clones containing ≥3 recovered cells are included. (D) Within-region comparisons of *Rbp1* expression between Mode 5-associated ECs and ECs assigned to other modes in the Trunk library (top; p=0.051), and *Ifitm1* expression between Mode 6-associated ECs and ECs assigned to other modes in the Tail library (bottom; p=0.00051). p-values were calculated using a two-sided Mann-Whitney U test. All panels show data from embryos barcoded at E6.5 (iE6.5).

**Figure S13:**
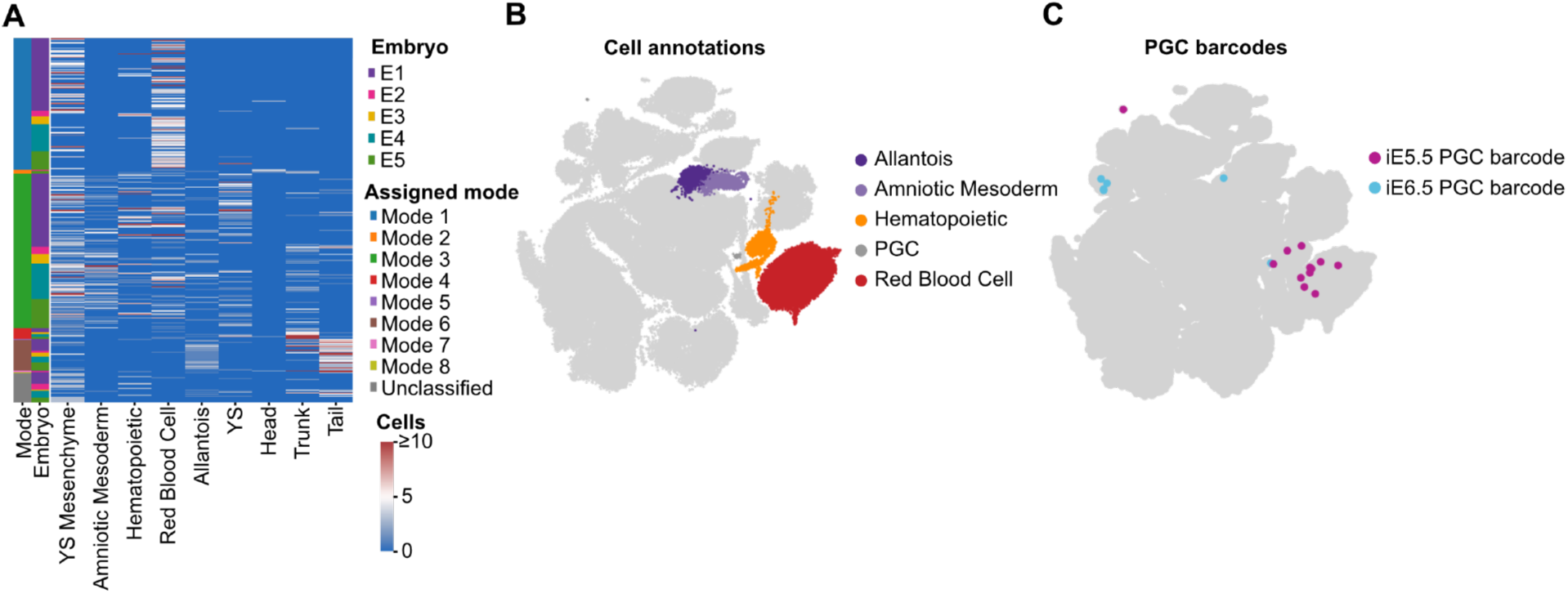
Supplemental data for extra-embryonic mesoderm origin analysis. (A) Heatmap showing the distribution of extraembryonic mesoderm (ExM)-containing clones across ExM and non-ExM populations from different anatomical regions. Rows represent unique barcodes, grouped and ordered by DMM mode assignment. Annotation bars indicate embryo identity and DMM mode. Color indicates the number of recovered cells, capped at 10. Only clones containing ≥3 recovered cells are included. (B) UMAP showing cell-type annotations for cells belonging to primordial germ cell (PGC)-containing clones. (C) Distribution of two PGC-containing clones across the transcriptomic UMAP, highlighted in blue and purple.

**Figure S14:**
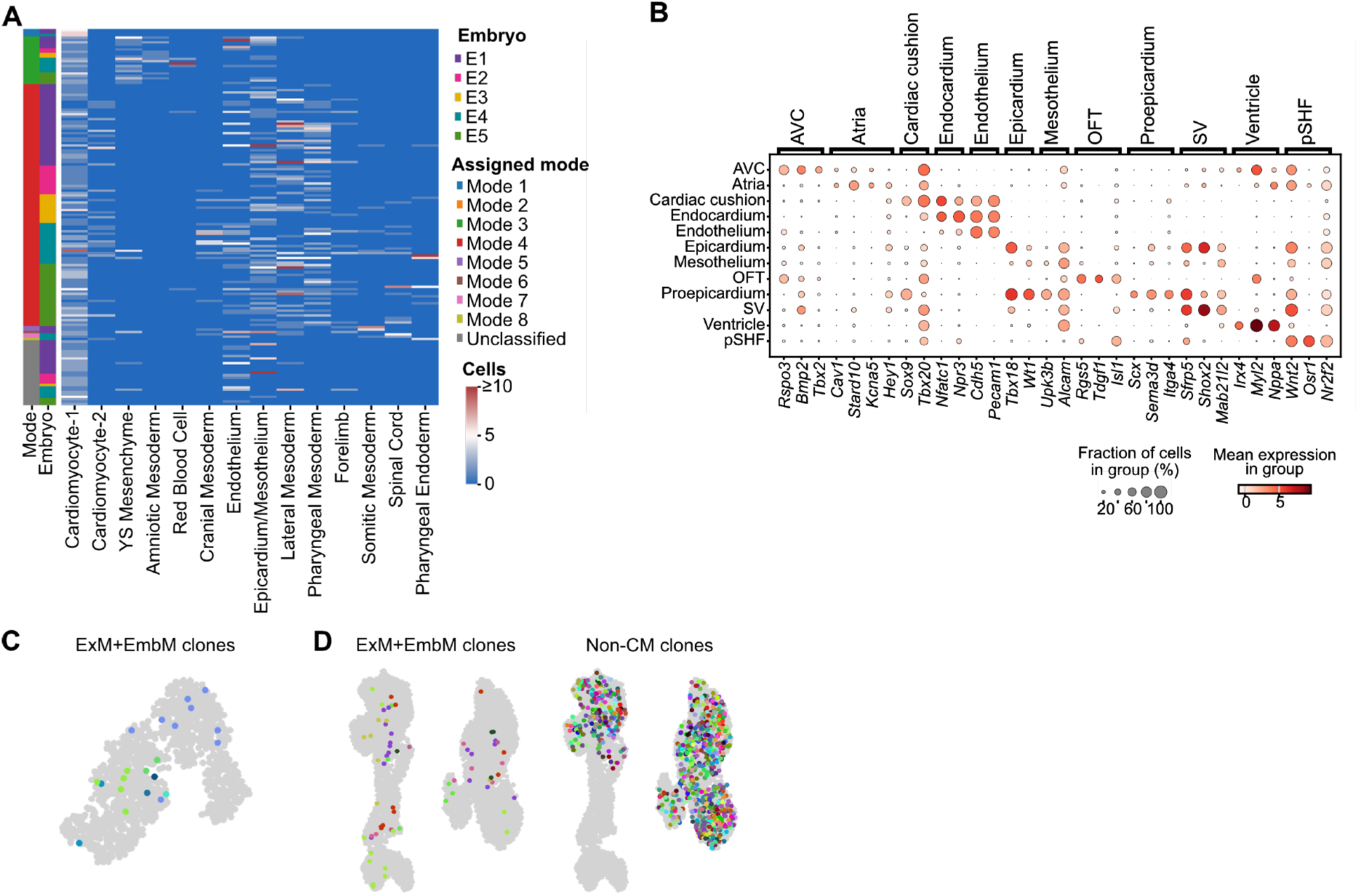
Supplemental data for cardiac cell origin analysis. (A) Heatmap showing the cell-type composition of cardiomyocyte-containing clones, grouped and ordered by DMM mode assignment and embryo. Rows represent unique barcodes and columns represent cell types. Annotation bars indicate embryo identity and DMM mode. Color indicates the number of recovered cells, capped at 10. Only clones containing ≥2 recovered cells are included. (B) Dot plot showing expression of marker genes used to annotate cardiac cell populations. (C) UMAP showing individual clones in “ExM+EmbM” categories, with each color representing a unique barcode. UMAP corresponds to those shown in 5B-D. (D) UMAP showing individual clones in “ExM+EmbM” and “Non-CM” categories, with each color representing a unique barcode. UMAP corresponds to those shown in 5F,G. All panels show data from embryos barcoded at E6.5 (iE6.5).

## Supplementary tables

**Table S1:** Summary of the total number of cells collected from each embryo and library and the number retained at each successive step of barcode processing and filtering.

**Table S2:** Results of differential gene expression analysis comparing endothelial cells across different anatomical regions of the embryo.

**Table S3:** Results of differential gene expression analysis comparing barcoded endothelial cells assigned to different DMM modes.

## Methods

### Mice

We used *Col1a1*^CARLIN^;*Tigre*^CARLIN^;*Rosa26*^CARLIN^ triple array mice (Jax strain #038750), *Col1a1*^Cas9^;*Rosa26*^M2^ mice (for CARLIN experiments; Jax strain #029415), and *Col1a1*^Cas9Dntt^Rosa26^M2^ mice (for DARLIN experiments; Jax strain #038749). Mice were maintained in homozygosity. For timed pregnancies, female mice were placed with male mice overnight and plugs were checked the following morning: noon on the day of observing the plug was designated E0.5. Mice were maintained in standard conditions of housing and husbandry at Boston Children’s Hospital or at Stanford University. All the procedures involving animals were approved by the Boston Children’s Hospital Institutional Animal Care and Use Committee and Stanford’s Administrative Panel on Laboratory Animal Care.

### Animal procedures

CARLIN/ Cas9 embryos were induced *in utero* with either 50μg/g or 25μg/g doxycycline (Dox, Sigma-Aldrich, catalog D9891) dissolved in PBS (10μg/ul concentration) via retro-orbital injection to the pregnant dam at E5.5 or E6.5. On the day of embryo collection, dams were euthanized via CO_2_ and the uterus was removed and placed in cold 2% FBS/PBS for embryo dissection.

### Isolation of tissues for single cell RNA-sequencing

Embryos were dissected in cold PBS + 2% FBS into head (cut above otic vesicle), trunk (otic vesicle to ∼8^th^ somite), tail and visceral YS (parietal YS was removed and discarded during dissection). For E1, the heart was microdissected and sequenced in a separate library to enrich for cardiac-related cell types. Each sample was incubated with TripLE Express Enzyme (ThermoFisher Scientific #12604013) at 37°C for 7 minutes. Tissues were further dissociated by gentle pipetting and the enzyme was blocked with cold FBS. The solution was passed through a 40µm strainer and spun down at 300g for 5 minutes at 4°C. Single cells resuspended in PBS and counted manually with a hemocytometer and Trypan blue staining. Encapsulation was performed using Chromium Single Cell 3′ Reagent Kits (v3.1 Dual Index chemistry for all embryos except the cardiac-enriched embryo (E5), which used v4 chemistry). A target of 10,000 cells was used for all embryos except the cardiac-enriched embryo, in which 20,000 cells were targeted.

### scRNA-seq library preparation and sequencing

Whole transcriptome libraries were generated following the protocol outlined by 10x Chromium and sequenced on NovaSeq (Illumina) through Novogene Corporation, aiming for 20,000 reads per cell. CARLIN/DARLIN barcode libraries were generated as previously described^49^. Briefly, following cDNA amplification, *Col1a1* and *Tigre* arrays were amplified through two nested PCR steps and indexed for sequencing. Kapa HiFi HotStart ReadyMix (Roche Sequencing Solutions, cat # 07958935001) was used for amplification and libraries were cleaned after each PCR step using SPRI beads (Beckman Coulter cat # B23318). Libraries were sequenced on Illumina MiSeq or MiSeq i100 plus using paired-end 500 cycles v2 kits or MiSeq i100 Plus 600 cycle kits with 10% PhiX sequencing control v3 (Illumina).

### Barcoding kinetics analysis

Timed pregnancies between *Col1a1*^CARLIN^ and *Col1a1*^Cas9^;*Rosa26*^M2^ mice were set up and embryos were induced with 25μg/g Dox on the 6^th^ day of pregnancy. At the indicated stage of collection (8,24,36,48,60h after Dox injection), embryos were dissected in cold PBS and placed in Trizol. Negative controls were uninduced and collected at E8.5-E9.5. RNA was extracted following manufacturer’s instructions, resuspended in water, and bulk CARLIN RNA sequencing libraries were generated as previously described. In brief, depending on the embryo stage, 20-100% of the total volume of RNA per embryo was loaded into a reverse transcription reaction with a CARLIN-specific primer, dNTPs (NEB), SuperScript III (Invitrogen), and RNAseOUT (Invitrogen). The cDNA was loaded into two nested PCR amplification reactions using the Q5 High Fidelity Polymerase (with purification using AMPure beads in between each stage) followed by an indexing reaction. Libraries were sequenced via Illumina MiSeq using paired-end 500 cycle v2 kits (Read 1: 250 cycles; Index Read: 6 cycles; Read 2: 250 cycles) with 10% PhiX sequencing control v3 (Illumina).

### Embryo immunofluorescence staining

Embryos were fixed in 4% PFA for 1h at 4°C on a shaker and washed 3 times in PBST (0.1% Tween-20) for 10 minutes each on a shaker at RT. Samples were permeabilized and blocked with 2% serum, 0.5% Triton in PBS for 1h on a shaker at RT. Embryos were stained overnight with Brachyury (cat number EPR18113; Abcam; 1:400) at 4°C in staining buffer (2% donkey serum PBST) and washed 3 times for 10 minutes each in PBST on a shaker. Secondary antibody (cat number A31573; Invitrogen; 1:1000) staining was performed overnight at 4°C in staining buffer (2% donkey serum PBST; 1:) and washed as described above. Samples were incubated with DAPI (1:1000) in PBST for 1h at 4°C in the dark and washed 3 times for 10 minutes each in PBST on a shaker. Samples were mounted in 10uL RapiClear (Sunjin lab #149001) on Superfrost slides with one 0.12mm Secure-Seal™ Spacer (Thermofisher # S24737). Embryos were imaged using an upright Zeiss LSM 980, 20x air objective.

### Data analysis

#### Whole transcriptome analysis pipeline

10x sequencing data were aligned to the mm10 reference genome using the standard 10x Genomics Cell Ranger pipeline using Cell Ranger v5.0.0 for all embryos except the cardiac-enriched embryo, which was processed using v9.0.1. The Vaeda filtering package^50^ (version 0.0.30) was used to filter out doublets using the default settings, and cells were filtered based on outlier mitochondrial/ gene counts. For the cardiac-enriched embryo, Vaeda filtering was performed post-gene/ mitochondrial filtering due to larger proportion of cell debris that resulted in inaccurate doublet estimations. We further excluded cells with fewer than 1,750-3,000 genes, high read counts, and more than 6-9% mitochondrial counts (depending on the dataset). Counts were log normalized and scaled to 10 standard deviations and datasets were integrated using Harmony on the PCA, with the embryo ID as a batch^51^. Harmonized PCAs were used for nearest neighbor graph calculation for UMAP visualization and Leiden clustering.

#### Cell annotations

We performed leiden clustering on the dataset and each cluster was assigned to a germ layer based on expression of markers shown in Figure S2F. To gain higher cell type resolution within each germ layer, we performed another round of leiden clustering on each individual germ layer dataset and annotated cells based on the expression of markers in Figure S2G-I. Cell type annotations were transferred back to cells in the combined dataset. Two Leiden clusters were removed based on outlier mitochondrial gene percentage and high ribosomal counts. Lateral plate mesoderm is denoted “Lateral Mesoderm” for conciseness.

#### CARLIN/DARLIN analysis pipeline

The CARLIN/DARLIN analysis was carried out as described before^19^. A cut-off of 10 reads per UMI and 10 reads per cell barcode was used. A template list of cell barcodes from the cell ranger output of the whole transcriptome library was used for read mapping.

#### Barcode filtering and merging

Barcodes were filtered out from the dataset if 1) they were observed in more than one embryo in our dataset, including two sequenced embryos not included in this manuscript, and 2) they contained a single edit. Barcode overlap was then calculated separately for CARLIN (E5.5 1, E3, E4, E5) and DARLIN (E5.5 2, E1, E2) embryos. First, Collagen and Tigre CARLIN barcode sequences were merged to create a unified clone ID for each cell. The Mosaic lineage package^20^ was used to collapse barcodes across cells using shared sequences at either CC or TC locus. In cases where only a subset of cells contained both Collagen and Tigre barcodes, these cells were used to link and merge larger groups of cells that shared only one locus and assigned them to a common clone as previously described^20^. Briefly the parameters used were:

- Unedited barcodes were given a significant p-value, though any barcodes composed of only unedited after running the pipeline were excluded.
- P-value threshold of 3×10^-5^ excludes unsignificant barcodes from the graph calculation.
- allele_N_cutoff=3. Up to 2 different alleles are allowed per clone (meaning 1 TC and 1 CC).
- Joint_allele_cutoff=6. Sequences with a joint allele above ≥6 are given the cutoff p-value. This downweights the pvalue of unedited barcodes so that it is not used to link clones that have different barcodes in another locus.
- Clones that were not combined by the pipeline retain their unified clone ID from the previous step unless they contain none. These barcodes were not merged because there were multiple sequences in one of the loci, therefore missing barcodes could be one of many sequences so they should not be connected and are excluded.

#### Clonal coupling analysis

##### Whole embryo coupling analysis

Our lineage-coupling analysis was based on the method described in Wagner et al. (2018)^23^. Extraembryonic endoderm was excluded because of negligible barcode editing (Figure S3B). We included clones containing at least three recovered cells and cell types represented in at least two clones. For each pair of cell types, the contribution of each clone containing both populations was calculated as the number of cells belonging to either cell type (k) divided by the total clone size (n). These contributions were summed across shared clones to obtain the observed lineage-coupling score (p).

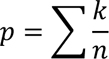

To determine whether the observed coupling exceeded that expected by chance given the sampling structure of the dataset, we generated 10,000 permutations in which cell-type labels were shuffled within each embryo, including for analyses of the pooled dataset. This preserved cell-type abundances, barcoded cell type frequencies and clone-size distributions within each embryo, while disrupting associations between cell-type and clone identities. Lineage coupling scores for each cell-type pair was standardized relative to its permutation-derived null distribution to obtain a Z-score, either across the pooled dataset (Figure 1C-D) or per individual embryo datasets (Figure S6-7). Coupling Z-scores were visualized as clustered heatmaps using hierarchical linkage clustering implemented in SciPy.

##### Coupling analyses of endothelial cells, extraembryonic mesoderm and allantois

To assess preferential lineage coupling between endothelial cells (ECs), extraembryonic mesoderm (ExM) or allantois and other cell populations, we performed conditional permutation analyses using the same lineage coupling score metric described above. Clone identity, clone size, focal cell-type identities and the number of non-focal cousins per clone were held fixed, while the joint cell-type x dissection-region labels of non-focal cells were shuffled in an embryo-stratified manner, preserving their embryo-specific abundances while disrupting their associations with individual clones. For ECs, non-EC cell-type and dissection-chunk identities were permuted together to assess preferential coupling between cell types from specific anatomical regions. The same approach was applied separately to ExM 1 (allantois) and ExM 2 (YS mesenchyme, amniotic mesoderm, blood) populations. For allantois, non-allantois cell-type identities were permuted to assess preferential coupling with individual cell types, irrespective of anatomical region.

Observed lineage-coupling scores were compared with permutation-derived null distributions to calculate Z-scores, quantifying enrichment or depletion of specific clonal partners conditional on the observed extent of non-focal sharing, rather than overall lineage restriction. Analyses were performed on pooled data and individual embryos, with embryo-specific null distributions calculated independently. Bootstrap resampling of clones was used to estimate 95% confidence intervals. For ECs and ExM, sensitivity analyses additionally permuted cell-type identities within each embryo × dissection chunk, preserving anatomical composition while testing cell-type-specific coupling.

### Concordance of lineage coupling between embryos

To assess the reproducibility of lineage-coupling patterns across embryos, we calculated coupling Z-scores for each embryo individually and compared these with the pooled Z-scores of all remaining embryos using Spearman correlation. To contextualize the resulting correlations, we generated “sampling-matched” and “scrambled” controls. For the sampling matched control, two disjoint sets of clones were drawn from the pooled dataset, matched in size to the individual embryo and the remaining embryos, respectively, and their coupling Z-scores were correlated (10 iterations, 1,000 permutations per iteration). For the scrambled control, cell-type labels were permuted within the individual embryo before calculating the correlation. Correlations were restricted to cell-type pairs represented in the corresponding individual embryo.

### Differential gene expression analysis

The PyDESeq2 package was used to perform the pseudo-bulk differential gene expression (DGE) analyses on our scRNA-seq dataset^52^. Only embryos with at least 5 cells in both groups were included in the analysis. DGE was calculated between ECs of a single mode with at least 3 embryos (Mode 3-6) and of all other modes combined (“all”), excluding unclassified. Similarly, DGE was calculated between ECs of one tissue with at least 3 embryos (Head, Trunk, Tail, YS) and of all other tissues combined (“all), including both barcoded and unbarcoded ECs. The heart library of was combined with the trunk library for E1. Raw counts were summed per embryo in each group. Only genes with at least 10 counts in at least 35% of groups (to ensure meaningful expression) were included in the analysis. Statistical significance was assessed with the default Wald Test. Genes were considered differentially expressed if they had an adjusted p-value (FDR) < 0.05 and an absolute log2 fold change > 1.0. To test for differences in expression levels of specific genes (*Rbp1* and *Ifitm1*) between modes in the same tissue (Trunk and Tail respectively), a two-sided Mann-Whitney U test was applied to the normalized counts of single ECs of that tissue grouped between Mode 5 and 6 respectively and all other modes combined (“other modes”), excluding unclassified cells.

### Modelling effective barcode recovery

To assess variation in barcode recovery across embryos and cell types, we fitted a beta-binomial model to the number of cells with usable barcodes relative to the total number of captured cells, stratified by embryo, sequencing library and cell type. The model estimated embryo-specific barcode recovery probabilities while allowing for additional variation beyond binomial sampling. Model fit was assessed using parametric simulations to generate predictive intervals for barcode recovery across embryos, anatomical libraries and cell types. Further details are provided in the Supplementary Statistical Methods document.

### Dirichlet-multinomial mixture modelling of clonal composition

To identify recurrent patterns of clonal contributions while accounting for incomplete sampling, we applied Dirichlet-multinomial mixture (DMM) modelling to the cell-type composition of individual clones. This approach models recovered cells as a finite sample from an underlying clonal composition and identifies recurrent compositional patterns, termed modes. Models containing 1-12 modes were fitted using the DirichletMultinomial package in R, with the optimal number of modes selected using the Laplace approximation to the model evidence. Clones were assigned to their most probable mode when its posterior probability exceeded twice that of the second-most probable mode; otherwise, they were classified as unclassified. Model robustness was evaluated through leave-one-embryo-out analyses and sensitivity analyses varying the minimum recovered clone size. Further details are provided in the Supplementary Statistical Methods.

### Pseudotime gene expression histograms

For plots S11D,E: Mean expression was computed on a uniform pseudotime grid using a sliding window encompassing 8% of cells, followed by Gaussian smoothing (σ = 2.5). Extreme pseudotime tails were capped (2nd-98th percentiles) to reduce endpoint artifacts, and grid positions supported by fewer than 30 cells were masked.

### Generative modelling of paraxial mesoderm clonal contributions

To assess whether the observed mesoderm-biased clonal contributions could be explained by incomplete sampling of mixed-lineage clones, we fitted four generative models accounting for population clone size, lineage composition and embryo-specific cell recovery (Supplementary Methods). Clone sizes were modelled using discretized log-normal distributions, allowing distinct mean sizes for different clonal populations. The models differed in whether they assumed a single population of mixed-lineage clones or an additional mesoderm-only population, and whether the mesodermal contribution of mixed clones was fixed or varied between clones according to a beta distribution. Specifically, M1 assumed a single mixed population with fixed lineage proportions; M2 additionally included a mesoderm-only population; M3 allowed heterogeneous lineage proportions within a single mixed population; and M4 combined heterogeneous mixed clones (beta distribution) with a mesoderm-only population.

Models were compared using predictive performance on held-out embryos, with 95% confidence intervals calculated by matched bootstrap resampling of clones within embryos. Model fit was additionally assessed through posterior predictive simulations comparing observed and simulated clone-size and lineage-composition distributions. Using the best-fitting model (M4), we estimated the probability that a sampled mesoderm cell belonged to a population clone containing at least 95% mesoderm cells, both across the dataset and for individual mesoderm subtypes. Further details are provided in the Supplementary Statistical Methods.

