## Supplemental Methods for "Whole embryo barcoding reveals convergent origins of mesodermal cells during mammalian gastrulation"

Secchi et al. 2026

### 1 Modelling effective barcode recovery across embryos

#### 1.1 Rationale

The probability that a captured single-cell RNA-sequencing cell yielded a usable clonal barcode was potentially different between experiments and embryos. Ignoring this variation would allow samples with different barcode-recovery efficiencies to contribute unequal amounts of clonal information and could confound technical and biological differences. We therefore modelled barcode recovery before analysing clone composition. We aimed to estimate reproducible embryo-level differences while allowing recovery fractions in small cell groups to vary more than expected under independent binomial sampling.

#### 1.2 Beta-binomial model

We aggregated cells into strata defined by embryo, sequencing library and cell-type annotation. This allowed us to test whether the observed variation in barcode recovery could be explained by embryo-specific differences and sampling variation alone.

For stratum $s$, let $n_s$ denote the number of captured annotated cells and $y_s$ the number with a usable barcode (see Methods).

We fitted

$$y_s \sim \mathrm{BetaBinomial}\left(n_s,\ \kappa\mu_s,\ \kappa(1-\mu_s)\right), \quad (1)$$

$$\mathrm{logit}(\mu_s) = \beta_0 + \beta_{e[s]}, \quad (2)$$

where $e[s]$ is the embryo containing stratum $s$, $\mu_s = \mu_{e[s]}$ is its embryo-specific effective recovery probability, and $\kappa$ is a concentration parameter shared across all strata.

Equivalently, the model allows a stratum-specific latent recovery probability

$$p_s \sim \mathrm{Beta}(\kappa\mu_{e[s]}, \kappa(1-\mu_{e[s]})) \quad (3)$$

$$y_s \mid p_s \sim \mathrm{Binomial}(n_s, p_s) \quad (4)$$

Its variance is

$$\mathrm{Var}(y_s) = n_s\mu_s(1-\mu_s)\frac{n_s+\kappa}{1+\kappa}. \quad (5)$$

Thus, large values of $\kappa$ approach independent binomial sampling, whereas smaller values model residual technical or biological heterogeneity.

We estimated all coefficients and $\kappa$ by a maximum likelihood approach, using L-BFGS-B optimisation with exact gradients calculated by JAX, three initialisations and bounds $\kappa \geq 0.02$.

#### 1.3 Model assessment

We assessed model adequacy by parametric simulation while retaining the observed number of cells in each stratum. For each fitted stratum and simulation,

$$p_s^{(b)} \sim \mathrm{Beta}(\widehat{\kappa}\widehat{\mu}_s, \widehat{\kappa}(1-\widehat{\mu}_s)), \quad (6)$$

$$y_s^{(b)} \sim \mathrm{Binomial}(n_s, p_s^{(b)}). \quad (7)$$

We used these simulations to obtain predictive intervals for recovery fractions aggregated by embryo, anatomical library and cell type.

### 2 Dirichlet–multinomial mixture analysis of sampled clonal composition

#### 2.1 Rationale: relating sampled cells to clonal behaviour

Analyses of clonal behaviour must distinguish uncertainty due to finite sampling from genuine differences between clones. A clone that produces several cell types in the embryo can appear restricted when only a few of its cells are recovered, whereas a larger sampled clone provides a more precise estimate of its composition.

Dirichlet–multinomial mixture models (DMMs) have been used to address an analogous problem in microbial community sequencing data [1]. Each microbial sample is represented by integer counts of multiple taxa, sequencing depth varies between samples and the underlying community compositions are themselves heterogeneous. Sequencing counts are modelled as a multinomial sample from an unobserved composition, while a Dirichlet distribution describes biological variation among compositions. A mixture of Dirichlet distributions thus represents recurrent compositional patterns.

We therefore represented each clone by an unobserved vector of cell-type proportions and treated its recovered cells as a finite multinomial sample from this vector. The Dirichlet layer allowed the underlying proportions to differ between clones, while the mixture model tested whether a single distribution adequately described this heterogeneity or whether it was better summarised by several recurrent distributions, which we refer to here as 'modes'.

This model is well suited to sparse clonal count data. Larger clones contribute more compositional information than smaller clones, and the Dirichlet distribution permits greater heterogeneity than expected under a fixed multinomial composition. In contrast to descriptive clustering of a barcode count matrix, the DMM provides an explicit probability model relating an underlying clonal composition to its sampled cell counts.

### 2.2 Cell and clone preprocessing

We filtered cells by those carrying a usable barcode, and complete cell-type and germ-layer annotations. We also excluded primordial germ cells, extraembryonic ectoderm and extraembryonic endoderm due to low sampling and deviations in barcode recovery rates.

For clone $i$, we aggregated cells into

$$\mathbf{y}_i = (y_{i1}, \dots, y_{iJ}), \qquad n_i = \sum_{j=1}^{J} y_{ij}, \tag{8}$$

where $y_{ij}$ is the number of sampled cells annotated as cell type $j$, and $n_i$ is the total number of sampled cells in that clone.

### 2.3 Generative model

For a model with $K$ modes,

$$z_i \sim \text{Categorical}(\pi_1, \dots, \pi_K), \tag{9}$$

$$\mathbf{p}_i \mid z_i = k \sim \text{Dirichlet}(\boldsymbol{\alpha}_k), \tag{10}$$

$$\mathbf{y}_i \mid \mathbf{p}_i, n_i \sim \text{Multinomial}(n_i, \mathbf{p}_i). \tag{11}$$

Here, $z_i$ is the unobserved mode of clone $i$, $\pi_k$ is the frequency of mode $k$ among the analysed clones, $\mathbf{p}_i$ is the latent composition of clone $i$, and $\boldsymbol{\alpha}_k = (\alpha_{k1}, \dots, \alpha_{kJ})$ defines the distribution of compositions within mode $k$. The mean composition and concentration of mode $k$ are

$$\theta_{kj} = \frac{\alpha_{kj}}{A_k}, \qquad A_k = \sum_{j=1}^{J} \alpha_{kj}. \tag{12}$$

The vector $\boldsymbol{\theta}_k$ gives the typical relative contribution of each cell type. Larger values of $A_k$ describe a comparatively homogeneous mode whose clones lie close to $\boldsymbol{\theta}_k$, whereas smaller values permit greater clone-to-clone heterogeneity, including clones dominated by different individual cell types.

Integrating over $\mathbf{p}_i$ gives the Dirichlet–multinomial probability

$$P(\mathbf{y}_i \mid z_i = k, n_i) = \frac{n_i!}{\prod_j y_{ij}!} \frac{\Gamma(A_k)}{\Gamma(A_k + n_i)} \prod_{j=1}^{J} \frac{\Gamma(\alpha_{kj} + y_{ij})}{\Gamma(\alpha_{kj})}, \tag{13}$$

and the likelihood for clone $i$ is

$$P(\mathbf{y}_i \mid n_i) = \sum_{k=1}^{K} \pi_k P(\mathbf{y}_i \mid z_i = k, n_i). \tag{14}$$

Because this probability is conditional on $n_i$, a one-cell clone can remain compatible with many underlying compositions, whereas a large clone with a distinctive count pattern provides stronger evidence.

### 2.4 Fitting and pooled choice of mode number

We fitted the DMM using `DirichletMultinomial::dmn` in Bioconductor [2]. We evaluated $K = 1, \dots, 12$. For each value of $K$, we initiated ten fits with different random seeds and retained the restart with the smallest Laplace criterion. Following Holmes *et al.*, we selected the primary pooled value of $K$ as the model with the smallest Laplace approximation to the negative log model evidence returned by `goodnessOfFit`. This criterion favours additional modes only when the improvement in fit outweighs greater model complexity.

### 2.5 Clone membership probabilities

For every clone, the fitted model gives

$$r_{ik} = P(z_i = k \mid \mathbf{y}_i, n_i) = \frac{\pi_k P(\mathbf{y}_i \mid z_i = k, n_i)}{\sum_{\ell=1}^{K} \pi_\ell P(\mathbf{y}_i \mid z_i = \ell, n_i)}. \tag{15}$$

We assigned each clone to its most probable mode only when the leading probability was more than twice the second-largest probability. We otherwise labelled the clone as unclassified.

#### 2.6 Leave-one-embryo-out sensitivity

As a sensitivity analysis, we performed leave-one-embryo-out refitting. We aligned modes from each leave-one-embryo-out fit one-to-one with the pooled modes using the Jensen–Shannon distance between their mean composition vectors and the Hungarian assignment algorithm.

We summarised agreement between fits $a$ and $b$ using the probability-weighted Jaccard index

$$J_k(a,b) = \frac{\sum_i \min(r_{ik}^{(a)}, r_{ik}^{(b)})}{\sum_i \max(r_{ik}^{(a)}, r_{ik}^{(b)})}. \tag{16}$$

#### 2.7 Sensitivity to sampled clone size

We repeated the $K = 1, \ldots, 12$ scan after imposing minimum sampled clone sizes of one to five cells. We then applied the calculations used for the leave-one-embryo-out sensitivity analysis.

### 3 Sampling-aware modelling of paraxial-mesoderm restriction

#### 3.1 Rationale and analysis population

Because we sampled only a fraction of barcoded cells, a clone that contributed both mesoderm and non-mesoderm cells in the embryo could appear mesoderm-only if none of its non-mesoderm descendants were sampled. We therefore used a generative model to test whether incomplete sampling alone could explain the observed clone counts or whether the data supported an additional mesoderm-restricted mode of clonal behaviour.

We included clones with at least one sampled Somitic Mesoderm or Presomitic Mesoderm cell from the Trunk or Tail, treating the Heart library from embryo PM655 as Trunk. We used $M$ to denote any cell annotated as mesoderm or extraembryonic mesoderm and $X$ to denote any retained non-mesoderm cell. We excluded primordial germ cells, extraembryonic ectoderm and extraembryonic endoderm because they were sampled at low frequency and had barcode-recovery rates that deviated from the embryo average (Section 1).

#### 3.2 Embryo-specific sampling

For each embryo, we first used the beta-binomial model described in Section 1 to estimate the probability $r_e$ that a captured annotated cell had a usable barcode. If $C_e$ is the number of captured eligible cells and $P_e$ is the total embryo cell population, the effective probability that a population cell contributed to the clonal dataset was

$$\pi_e = r_e \frac{C_e}{P_e}. \tag{17}$$

We applied the same $\pi_e$ to mesoderm and non-mesoderm cells within an embryo. We represented uncertainty in embryo size by a normal distribution centred on 500,000 cells, for which 300,000–700,000 cells corresponded to its 95% interval. We propagated draws from the recovery and population-size distributions through the bootstrap analysis. Maximum-likelihood fits used the median $\pi_e$ for each embryo.

#### 3.3 Model

For clone $i$, let $N_i$ denote its total population size and $\theta_i$ its population fraction of non-mesoderm cells. Models containing a distinct mesoderm-only class assigned

$$P(Z_i = M\text{-only}) = q, \qquad P(Z_i = \text{mixed}) = 1 - q, \tag{18}$$

whereas models without this class fixed $q = 0$. Mesoderm-only clones had $\theta_i = 0$. For mixed clones, we either fixed the non-mesoderm fraction across clones or allowed it to vary:

$$\theta_i = \theta \quad \text{or} \quad \theta_i \sim \mathrm{Beta}(\alpha, \beta). \tag{19}$$

The Beta model therefore allowed a heterogeneous range of mesoderm biases within the mixed-clone category.

We modelled population clone size using a discretised log-normal distribution. In models containing a mesoderm-only class, we allowed mesoderm-only and mixed clones to have different mean sizes, $\mu_M$ and $\mu_{ME}$, but used a shared log-scale standard deviation $\sigma_N$. We applied the same embryo-level shift to both means,

$$\delta_e \sim \mathcal{N}(0, \tau_\delta^2), \tag{20}$$

which allowed the overall clone-size scale to vary between embryos, accounting for differences in sampling or labelling frequencies, without changing the relative size difference between clone classes.

Given $N_i$, $\theta_i$ and embryo $e$, we modelled each population cell as observed with probability $\pi_e$. The likelihood jointly described the observed numbers of non-mesoderm cells, paraxial-mesoderm anchor cells, other mesoderm cells and unobserved cells, conditional on observing at least one paraxial-mesoderm anchor cell.

#### 3.4 Four fitted hypotheses

M1 tested whether a common mixed behaviour combined with incomplete sampling was sufficient. M2 added continuous variation in mixed-clone composition. M3 instead added a distinct mesoderm-only class, whereas M4 included both forms of heterogeneity.

#### 3.5 Fitting, validation and sensitivity

We estimated parameters by maximum likelihood. We numerically integrated the likelihood over population clone size, mixed-clone composition and the shared embryo-level clone-size shift. All clones from the same embryo informed a common latent $\delta_e$.

We assessed replication by repeating the following procedure for each of the five embryos. We withheld one embryo, fitted each candidate model to clones from the remaining four embryos and used the fitted model to predict the observed mesoderm and non-mesoderm counts of every clone in the withheld embryo. We integrated uncertainty in its shift over the distribution learned from the training embryos. We summarised predictive performance as the mean log probability assigned to a held-out clone. To compare two models, we calculated the difference between their log probabilities for the same held-out clones. We resampled clones with replacement within each held-out embryo to obtain uncertainty intervals and averaged the five embryo-specific differences with equal weight, preventing embryos with more detected clones from dominating the comparison.

We estimated uncertainty in parameters and derived population summaries from 1,000 bootstrap fits. Within each embryo, we resampled clones and paired each bootstrap replicate with one draw from the barcode-recovery and embryo-size distributions before refitting all four models. Reported 95% intervals are bootstrap percentile intervals.

We assessed model adequacy by simulating datasets from each fitted model using embryo-specific sampling probabilities. We compared the observed and simulated distributions of detected clone sizes, the persistence of clones with no sampled non-mesoderm cells as detected clone size increased, and the composition of detected mixed clones.

#### 3.6 Calculating mesoderm restriction by clone

A distinct mesoderm-only class and a strongly mesoderm-enriched continuous distribution are biologically similar: under either model, most clones in the population are completely or almost completely mesodermal. We therefore operationally defined a clone as mesoderm-restricted if

$$\frac{M_i^{\mathrm{pop}}}{N_i} \geq 0.95. \tag{21}$$

From this, we calculated

$$R_i = P(M_i^{\mathrm{pop}}/N_i \geq 0.95 \mid X_i, M_i, A_i, e_i, \text{model}) \,, \tag{22}$$

where $M_i$ is the total sampled number of mesoderm and extraembryonic-mesoderm cells in clone $i$, while $A_i \subseteq M_i$ is the number of sampled Trunk or Tail Somitic Mesoderm or Presomitic Mesoderm cells used to determine clone eligibility. Thus, $0 < A_i \leq M_i$, and the sampled count of other mesoderm cells is $M_i - A_i$. $X_i$ is the sampled non-mesoderm count. This probability integrated over population size, composition, clone class and the embryo shift. To avoid fitting and scoring the same embryo, we calculated the primary $R_i$ values for each embryo using the model fitted to the other four embryos.

We used the continuous clone-level probabilities $R_i$ in two distinct downstream summaries. First, we summarised mesoderm restriction within each paraxial-mesoderm subtype or anatomical region as either the mean $R_i$ across contributing clones or the mean $R_i$ across sampled paraxial-mesoderm cells. Second,

| Model | Explicit $M$-only class | Mixed composition | Clone-size means |
|---|---|---|---|
| M1 | absent ($q = 0$) | fixed $\theta$ | $\mu_{ME}$ |
| M2 | absent ($q = 0$) | $\theta_i \sim \mathrm{Beta}(\alpha, \beta)$ | $\mu_{ME}$ |
| M3 | present ($q$ fitted) | fixed $\theta$ | $\mu_M, \mu_{ME}$ |
| M4 | present ($q$ fitted) | $\theta_i \sim \mathrm{Beta}(\alpha, \beta)$ | $\mu_M, \mu_{ME}$ |

among the probability-weighted non-restricted component of the data, we quantified the relative representation of each non-mesoderm partner group using $L_g$, as defined below.

For clone $i$, let $R_i$ denote the model-estimated probability that its population clone is at least 95% mesoderm, $n_{ig}$ the sampled number of non-mesoderm cells belonging to group $g$, and $n_{i,nonM}$ the total sampled number of non-mesoderm cells. We calculated linkage to group $g$ as

$$L_g = \frac{\sum_i (1 - R_i) n_{ig}}{\sum_i (1 - R_i) n_{i,nonM}} \tag{23}$$

[2] Morgan M. *DirichletMultinomial: Dirichlet–multinomial mixture model machine learning for microbiome data*. Bioconductor package. doi:10.18129/B9.bioc.DirichletMultinomial.
